# Genome-wide analysis in *Escherichia coli* unravels an unprecedented level of genetic homoplasy associated with cefotaxime resistance

**DOI:** 10.1101/2020.06.01.128843

**Authors:** Jordy P.M. Coolen, Evert P.M. den Drijver, Jaco J. Verweij, Jodie A. Schildkraut, Kornelia Neveling, Willem J.G. Melchers, Eva Kolwijck, Heiman F.L. Wertheim, Jan A.J.W. Kluytmans, Martijn A. Huynen

**Affiliations:** Department of Medical Microbiology and Radboudumc Center for Infectious Diseases, Radboud University Medical Center, Nijmegen, The Netherlands; Department of Infection Control, Amphia Ziekenhuis, Breda, The Netherlands; Laboratory for Medical Microbiology and Immunology, Elisabeth-Tweesteden Hospital, Tilburg, The Netherlands; Department of Human Genetics, Radboud University Medical Center, Nijmegen, The Netherlands; Laboratory for Microbiology, Microvida, Location Breda, The Netherlands; Julius Center for Health Sciences and Primary Care, UMCU, Utrecht, The Netherlands; Centre for Molecular and Biomolecular Informatics, Radboud University Medical Center, Nijmegen, The Netherlands

**Keywords:** *Escherichia Coli*, Genomics, Whole genome sequencing, *ampC*, Bioinformatics

## Abstract

Cefotaxime (CTX) is a commonly used third-generation cephalosporin (3GC) to treat infections caused by *Escherichia coli*. Two genetic mechanisms have been associated with 3GC resistance in *E. coli*. The first is the conjugative transfer of a plasmid harboring antibiotic resistance genes. The second is the introduction of mutations in the promoter region of the *ampC* β-lactamase gene that cause chromosomal-encoded β-lactamase hyperproduction. A wide variety of promoter mutations related to AmpC hyperproduction have been described. However, their link to a specific 3GC such as CTX resistance has not been reported. Here, we measured CTX MICs in 172 cefoxitin resistant *E. coli* isolates and performed genome-wide analysis of homoplastic mutations associated with CTX resistance by comparing Illumina whole-genome sequencing data of all isolates to a PacBio tailored-made reference chromosome. We mapped the mutations on the reference chromosome and determined their occurrence in the phylogeny, revealing extreme homoplasy at the −42 position of the *ampC* promoter. The 24 occurrences of a “T” at the −42 position rather than the wild type “C”, resulted from 18 independent C>T mutations in 5 phylogroups. The −42 C>T mutation was only observed in *E. coli* lacking a plasmid-encoded *ampC* gene. The association of the −42 C>T mutation with CTX resistance was confirmed to be significant (FDR < 0.05). To conclude, genome-wide analysis of homoplasy in combination with CTX resistance identifies the −42 C>T mutation of the *ampC* promotor as significantly associated with CTX resistance and underline the role of recurrent mutations in the spread of antibiotics resistance.

**Impact Statement:** In the past decades, the worldwide spread of extended spectrum beta-lactamases (ESBLs) has led to a substantial increase in the prevalence of resistant common pathogens, thereby restricting available treatment options. Although acquired resistance genes, e.g. ESBLs, get most attention, chromosome-encoded resistance mechanisms may play an important role as well. In E. coli chromosome-encoded β-lactam resistance can be caused by alterations in the promoter region of the *ampC* gene. To improve our understanding of how frequently these alterations occur, a comprehensive interpretation of the evolution of these mutations is essential. This study is the first to apply genome-wide homoplasy analysis to better perceive adaptation of the E. coli genome to antibiotics. Thereby, this study grants insights into how chromosomal-encoded antibiotic resistance evolves and, by combining genome-wide association studies with homoplasy analyses, provides potential strategies for future association studies into the causes of antibiotics resistance.

**Data summary:** All data is available under BioProject: PRJNA592140. Raw Illumina sequencing data and metadata of all 171 *E. coli* isolates used in this study is available from the Sequence Read Archive database under accession no. SAMN15052485 to SAMN15052655. Full reference chromosome of ampC_0069 is available via GenBank accession no. CP046396.1 and NCBI Reference Sequence: NZ_CP046396.1.

## Introduction

*Escherichia coli* is an important pathogen in both community and healthcare-associated infections [1,2]. In the past decades, a substantial increase in resistance to third-generation cephalosporin (3GC) antibiotics in *E. coli* has been observed worldwide, mainly caused by the production of extended-spectrum β-lactamases (ESBL) and AmpC β-lactamases, restricting available treatment options for common infections [3]. AmpC β-lactamases differ from ESBL as they not only hydrolyze broad-spectrum penicillins and cephalosporins, but also cephamycins. Moreover, AmpC β-lactamases are not inhibited by ESBL-inhibitors like clavulanic acid [3], limiting antibiotic treatment options even further. A widely used screening method for AmpC production is the use of cefoxitin (FOX) susceptibility, a member of the cephamycins [4].

Although *ampC* β-lactamase genes can be plasmid-encoded (*pampC*), they are also encoded on the chromosomes of numerous Enterobacterales. *E. coli* naturally carries a chromosomal-mediated *ampC (campC*) gene, but unlike most other Enterobacterales this gene is non-inducible due to the absence of the *ampR* regulator gene [3]. Consequently, chromosomal AmpC production in *E. coli* is exclusively regulated by promoter and attenuator mechanisms. This results in constitutive low-level *campC* expression that still allows the use of 3GC antibiotics, such as cefotaxime (CTX), to treat *E. coli* infections [3]. However, various mutations in the promoter/attenuator region of *E. coli* may cause constitutive hyperexpression of *campC* [5,6], thereby increasing the Minimal Inhibitory Concentrations (MICs) for broad-spectrum penicillins and cephalosporins and limiting appropriate treatment options.

A wide variety of promoter and attenuator mutations have been related to AmpC hyperproduction [6]. AmpC hyperproduction is primarily caused by alterations of the *ampC* promoter region, leading to a promoter sequence that more closely resembles the *E. coli* consensus sigma 70 promoter with a TTGACA −35 box separated by 17 bp from a TATAAT −10 box. These alterations can be divided into different variants associated with e.g. an alternate displaced promoter box, a promotor box mutation or an alternate spacer length due to insertions [6]. Furthermore, mutations of the attenuator sequence can lead to changes in the hairpin structure that strengthen the effect of promoter alterations on AmpC hyperproduction. In a study on cefoxitin-resistant *E. coli* isolated from Canadian hospitals, Tracz *et al*. described 52 variants of the promoter and attenuator region [6]. In this study a two-step quantitative reverse transcriptase (qRT-)PCR was used to determine the effect of promoter/attenuator variants on *ampC* expression. Various mutations were related to different delta–delta cycle threshold values in the RT-PCR and corresponding variations in FOX resistance. An interesting observation that emerged from this study was that the −32T>A and the −42C>T mutation were the major alterations that strengthened the *ampC* promoter. Both result in a consensus −35 box. Although it is known that AmpC hyperproduction leads to FOX resistance as studied by Tracz *et al*., the effect of various mutations on resistance to a 3GC antibiotic such as CTX have not been explored. This is relevant because CTX is commonly used in the treatment of patients with severe *E. coli* infections, often in combination with selective digestive tract decontamination in Intensive Care Units [7,8].

While previous research mainly focused on the chromosomal AmpC resistance mechanism and the impact of AmpC hyperproduction, there is a lack in knowledge and understanding of the evolutionary origin of these promoter/attenuator variants. Notably, it is unexplored how the two most prominent promoter mutations −32T>A and −42C>T are distributed over the *E. coli* phylogeny and therewith how often they occur. More precisely, literature shows selective pressure can lead to convergent evolution that results in the reoccurrence of a mutation in multiple isolates independently and in separate lineages [9]. This phenomenon is named homoplasy [10]. A Consistency Index can be calculated to quantify homoplasy by dividing the minimum number of changes on the phylogeny by the number of different nucleotides observed at that site minus one [11], effectively quantifying how often the same mutation occurred in a phylogenetic tree. One can use the Consistency Index to recognize genomic locations subjected to homoplasy, and relate the single-nucleotide polymorphism (SNP) positions that are inconsistent with the phylogeny to antibiotic resistance, as has e.g. been done in multiple studies on *Mycobacteria* spp [12–15].

In the present study, we hypothesize that some of the mutations in the *ampC* promoter/attenuator region are homoplastic and are associated with CTX resistance. To test our hypothesis, we performed genome-wide homoplasy analysis and combined it with a genome-wide analysis of polymorphisms associated with CTX resistance by constructing a tailored *E. coli* reference chromosome and combining it with WGS data of 172 FOX resistant *E. coli* isolates previously collected by our research group [16].

## Methods

### Isolate selection, DNA isolation, library preparation and DNA sequencing

One hundred seventy-two *Escherichia coli* isolates previously used by our study group [16] were selected in the present study (see Table S1 in supplemental material). To summarize the method; DNA isolation was performed as previously described [16], library preparations were performed using Illumina Nextera XT library preparation kit (Illumina, San Diego, CA, USA), and DNA sequencing was performed using an Illumina NextSeq 500 (Illumina, San Diego, CA, USA) to generate 2x 150bp paired-end reads or 2x 300 bp reads on an Illumina MiSeq (Illumina, San Diego, CA, USA). *De novo* assembly was also performed identical to the method as described in Coolen *et al*. 2019 [16] using SPAdes version 3.11.1 [17].

### Phylogroup and MLST

Phylogroup stratification was performed using ClermonTyping version 1.4.0 [18]. MLST STs were derived from the contigs using mlst version 2.5 pubMLST, 31 October 2017 [19,20].

### Obtaining the *ampC* promoter/attenuator region

To detect the promoter/attenuator region a custom blast database [21] was created using the 271 bp fragment as described by Peter-Getzlaff *et al*.[22] using *Escherichia coli* K-12 strain ER3413 (accession: CP009789.1) ABRicate version 0.8.9 [23] was used to locate matching regions per sample and were extracted and converted into multi-fasta format using a custom python script. Strains were labelled AmpC hyperproducer when promoter mutations were found, as reported by Caroff *et al*. [24] and Tracz *et al*. [6].

### Plasmid-mediated ampC detection

Detection of *pampC* genes was performed by using ABRicate version 0.5 [23] and ResFinder database (2018-02-16) as described by Coolen *et al*. [16].

### PacBio single molecule real-time (SMRT) sequencing of *E. coli* isolate

For PacBio SMRT sequencing, genomic DNA (gDNA) was extracted using the Bacterial gDNA Isolation Kit (Norgen Biotek Corp., CAN, ON, Thorold). A single *E. coli* isolate was subjected for DNA shearing using Covaris g-TUBEs (Covaris Inc, US, MA, Woburn) for 30 seconds on 11,000 RPM (*g*). Each DNA sample was separated into two aliquots. Size selection was performed using a 0.75% agarose cassette and marker S1 on the BluePippin (Sage Science Inc, US, MA, Beverly) to obtain either 4-8 kb or 4-12 kb DNA fragments. This size selection was chosen to maintain all DNA fragments including these originating from plasmids (data not used in this study). Library preparation was performed using the SMRTbell Template prep kit 1.0 (Pacific Biosciences, US, CA, Menlo Park). For cost-effectiveness, samples were barcoded and pooled with other samples that are not relevant for this study. Sequencing was conducted using the PacBio Sequel I (Pacific Biosciences, US, CA, Menlo Park) on a Sequel SMRT Cell 1M v2 (Pacific Biosciences, US, CA, Menlo Park) with a movie time of 10 h (and 186 min pre-extension time). Subreads per sample were obtained by extracting the bam files using SMRT Link version 5.1.0.26412 (Pacific Biosciences, US, CA, Menlo Park).

### Chromosomal reconstruction using *de novo* hybrid assembly

To obtain a full-length chromosome, Unicycler version 0.4.7 [25] (settings: --mode bold) was used, combining Illumina NextSeq 500 2x 150 bp paired-end reads with PacBio Sequel SMRT subreads. Because unicycler requires fasta reads as input, the subreads in bam format were converted to fasta by using bam2fasta version 1.1.1 from pbbioconda (https://github.com/PacificBiosciences/pbbioconda) prior to *de novo* hybrid assembly. The full circular chromosome was uploaded to NCBI and annotated using the NCBI Prokaryotic Genome Annotation Pipeline (PGAP) version 4.10 [26,27].

### SNP analysis using *E. coli* reference ampC_0069

Alignment of Illumina reads and SNP calling was performed for all isolates to the reference chromosome of *E. coli* isolate ampC_0069 using Snippy version 4.3.6 (https://github.com/tseemann/snippy). A full-length alignment (fullSNP) and a coreSNP alignment containing SNP positions shared among all isolates were generated by using snippy-core version 4.3.6 (https://github.com/tseemann/snippy).

### Inferring of phylogeny

A phylogenetic tree was inferred by using the coreSNP alignment as input for FastTree(MP) version 2.1.3 SSE3 (settings: -nt -gtr) [28].

### Detection of Homoplasy

The Consistency Index for all nucleotide positions on the chromosome was calculated using HomoplasyFinder version 0.0.0.9000 [10]. The coreSNP phylogeny was used as true phylogeny and the Consistency Index was calculated using the multiple sequence alignments fullSNP alignment as input.

### Relate mutations to CTX resistance

To assess if certain mutations were linked to CTX resistance all non-plasmid harboring *ampC E. coli* isolates were used. CTX resistance was defined using EUCAST guidelines standards of CTX MIC > 2 mg/L [29]. CTX MIC results were obtained from our previous study [16]. For each nucleotide position on the reference chromosome the number of Resistant and Sensitive isolates were counted and tested for adenine vs all other nucleotides, thymine vs all other nucleotides, cytosine vs all other nucleotides, and guanine vs all other nucleotides creating a contingency table and performing a Fisher Exact Test in R 3.6.1 [30]. To correct for multiple testing, *P* values were adjusted using FDR [31].

### Selection of genomic positions of interest

By combining previous metrics most relevant genomic positions were selected. Criteria for selection are, FDR <= 0.05 to CTX and Consistency Index of <= 0.05882353. Annotation of mutation positions was obtained by using the genome annotation of reference chromosome (GenBank accession no. CP046396) and applying snpEff (version 4.3t) [32]. The EnteroBase core-genome MLST and whole-genome MLST schemes were used to distinguish core and accessory genes [33].

### Recombination analysis

Gubbins version 2.4.1. (settings: -f 30) was used to detect recombination regions with coreSNP alignment and tree as input [34].

### Visualization of data

The interactive Tree of Life web-based tool iTOL version 5.3 was used to visualize the phylogenetic tree [35]. Information about CTX resistance, presence of the *pampC* gene, *campC* hyperproduction as defined, MLST and phylogroup, as well as alignments of promoter and attenuator region were incorporated into visualization. The sequence logo of the promoter and the attenuator alignment were generated using the web-based application Weblogo version 3.7 [36] (http://weblogo.threeplusone.com). A chromosome ideogram of the *E. coli* isolate ampC_0069 reference chromosome was visualized using CIRCOS software package version 0.69-8 [37]. Consistency Index scores and significant mutations associated with CTX resistance were plotted in the ideogram. Gubbins results were displayed by using phandango [38].

### Overview of method

A workflow graph of the methods is visualized in Fig 1 using the web-based application yEd Live version 4.4.2 (https://www.yworks.com/yed-live/).

**FIG 1.**
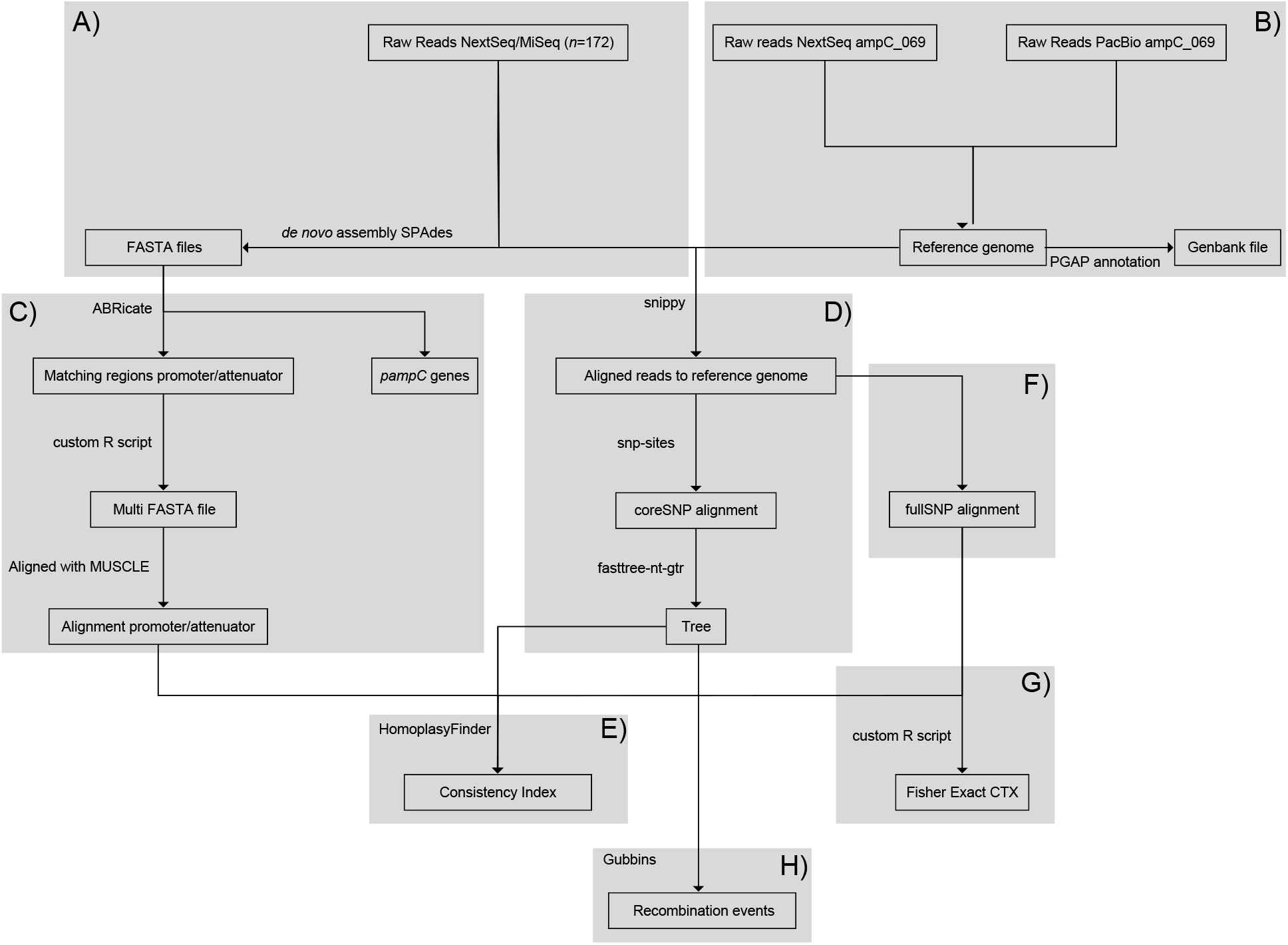
Schematic of workflow used to perform the homoplasy-based association analysis. Starting from the top A) the *de novo* assembly of the NextSeq/MiSeq reads and B) the hybrid assembly of the reference chromosome ampC_069. On the left side C) the alignment of promoter/attenuator region. In the middle D) the coreSNP analysis for the phylogeny used in E) the homoplasy analysis combined with F) the fullSNP data on the right, which was also used for G) the statistics (Fisher Exact & FDR) to relate cefotaxime (CTX) resistance to SNP positions. H) Inferring recombination events using Gubbins.

## RESULTS

### *E. coli* collection

To study genetic homoplasy events in suspected AmpC producing *Escherichia coli*, FOX MIC > 8 ug/ml and ESBL phenotype negative *E. coli* isolates (*n*=172) were selected as previously described by Coolen *et al*. [16] (see Table S1 in supplemental material). The entire collection was subjected to whole-genome sequencing followed by *de novo* assembly of the sequence reads to obtain contigs.

### MLST and phylogroup variants

To access the genetic diversity of our *E. coli* collection, we identified both multi-locus sequence typing (MLST) and phylogroup variants of each of the 172 *E. coli* isolates. Seventy-five different sequence types (STs) were identified, of which ST 131 (8.1% *n*=14), ST 38 (7.0% *n*=12), and ST 73 (7.0% *n*=12) were the most prevalent. The sequence types of 13 isolates are unknown. Phylogroup stratification revealed that the isolates belonged to eight different phylogroups (Table 1). Phylogroup B1, B2, and D were the most prevalent. One isolate belonged to *Escherichia* clade IV (st. no. ampC_0128).

**TABLE 1.** Table of the distribution of AmpC promoter and attenuator variants as well as the amount of different MLST and phylogroups per grouped genotype (*pampC*, hyperproducers and low-level AmpC producers).

|  | Isolates | Promoter variants | Attenuator variants | MLST | Phylogroups |
| --- | --- | --- | --- | --- | --- |
| *pampC* | *n*=90 | *n*=3 | *n*=6 | 44 STs & 4 unknown | A (11.1%), B1 (13.3%), B2 (27.8%), C (2.2%), D (31.1%), E (3.3%), F (11.1%) |
| Hyperproducers | *n*=59 | *n*=12 | *n*=14 | 30 STs & 5 unknown | A (1.7%), B1 (30.5%), B2 (50.9%), C (8.5%), D (5.1%), F (1.7%), clade IV (1.7%) |
| Low-level AmpC producers | *n*=23 | *n*=10 | *n*=5 | 14 STs & 4 unknown | A (4.4%), B1 (13.0%), B2 (39.1%), C (8.7%), D (30.4%), E (4.4%) |
| Total | *n*=172 | *n*=22 | *n*=18 | 75 STs and 13 unknown | A (7.0%), B1 (19.2%), B2 (37.2%), C (5.2%), D (22.1%), E (2.3%), F (6.4%), clade IV (0.6%) |

### *ampC* promoter and attenuator variants

We examined the whole *E. coli* genome. However, we firstly focused on mutations in the *ampC* promoter and attenuator region. Previously described mutations in the *ampC* promoter region that according to Tracz *et al*. lead to “hyperproduction” of AmpC were detected in 59 (34.4%) of the isolates [6]. These isolates were therefore labelled as hyperproducers. Analysis of the promotor area (−42 to −8) resulted in 21 different variants and the wild type (see Table 1). In the attenuator region (+17 to +37), 18 different variants were identified (see Table 1). One isolate (ampC_0128) showed an unusual promoter variant, a four-nucleotide deletion (−45_-42delATCC). Moreover, an insertion (21_22insG) of unknown function was detected in the attenuator (see Table S1 in supplemental material).

### Plasmid-mediated *ampC* variants

As we aim to associate chromosomal mutations with CTX resistance, differentiation of *pampC* harboring isolates from non-*pampC* harboring isolates was required. Genomic analysis showed that 90 (52.3%) of the isolates harbored a *pampC* gene of which *bla*_CMY-2_ was the most prevalent (*n*=78). One isolate harbored two different *pampC* genes (*bla*_CMY-4_ and *bla*_DHA-1_) (ampC_0119). One isolate contained a *bla*_CTX-M-27_ gene combined with a *bla*_CMY-2_ gene (ampC_0114) but was ESBL disc test negative (see Table S1 in supplemental material). In 23 (13.4%) of the isolates neither *pampC* nor described mutations related to AmpC hyperproduction were detected and are noted as low-level AmpC producers.

### Tailored reference chromosome

To be able to reconstruct an accurate phylogeny we selected *E. coli* isolate ampC_0069, one of the strains of the study, to use as reference chromosome for SNP calling. The tailored reference chromosome was constructed through a hybrid assembly of *n*=4,423,109 2x 150 bp Illumina NextSeq 500 paired-end reads together with *n*=218,475 PacBio Sequel SMRT subreads (median 5,640 bp). This resulted in a high-quality full circular chromosome of *E. coli* isolate ampC_0069, with a size of 5,056,572 bp. This isolate belongs to ST648 and contains a plasmid-encoded *bla*_CMY-42_. The full circular chromosome was uploaded to GenBank accession no. CP046396 and was used for further analysis. Genome annotation with the NCBI Prokaryotic Genome Annotation Pipeline (PGAP) identified 4,720 Coding Sequences.

### SNP calling

To be able to reconstruct the phylogeny and obtain SNP positions, we mapped reads of all isolates to the reference chromosome *E. coli* ampC_0069, (accession no. CP046396) resulting in a coreSNP alignment containing 314,200 variable core SNP positions. For further details per isolate see Table S2 in supplemental material. To validate our SNP calling method we compared the ampC_0069 Illumina NextSeq 500 paired-end reads to the reference chromosome of ampC_0069, resulting in 0 SNPs detected, supporting that the SNP calling data and method produce no false positives.

### Phylogenetic tree based on coreSNP

The coreSNP alignment was used for further analysis. Figure 2 illustrates the approximately maximum-likelihood phylogenetic tree of all 172 isolates based on the coreSNP alignment. The tree has a robust topology as indicated by computerized adaptive testing (CAT) likelihood calculations, resulting in only three positions with a value ≤60% [28]. When focusing on the *ampC* promoter mutations, they were most prevalent in phylogroups B1, B2, and C, although they were present in all phylogroups except phylogroup E that lacked mutations in either the promoter or attenuator region. Interestingly, two positions previously highlighted by Tracz *et al*., −42 and −32, are only mutated in the absence of a *pampC* gene, even in isolates with a similar MLST (ST12, ST88, and ST131). The −42C>T mutation, which results in an alternate displaced promoter box and therefore leads to increased resistance [6], is present in 24 isolates in 5 distinct phylogroups and in 17 separate phylogenetic branches, indicating that this mutation is homoplastic. Additionally, the −32T>A mutation in the promoter, previously also associated to resistance [6], is present in 20 isolates in 3 distinct phylogroups and in 14 separate phylogenetic branches. To quantify the level of homoplasy we calculated the Consistency Index.

**FIG 2.**
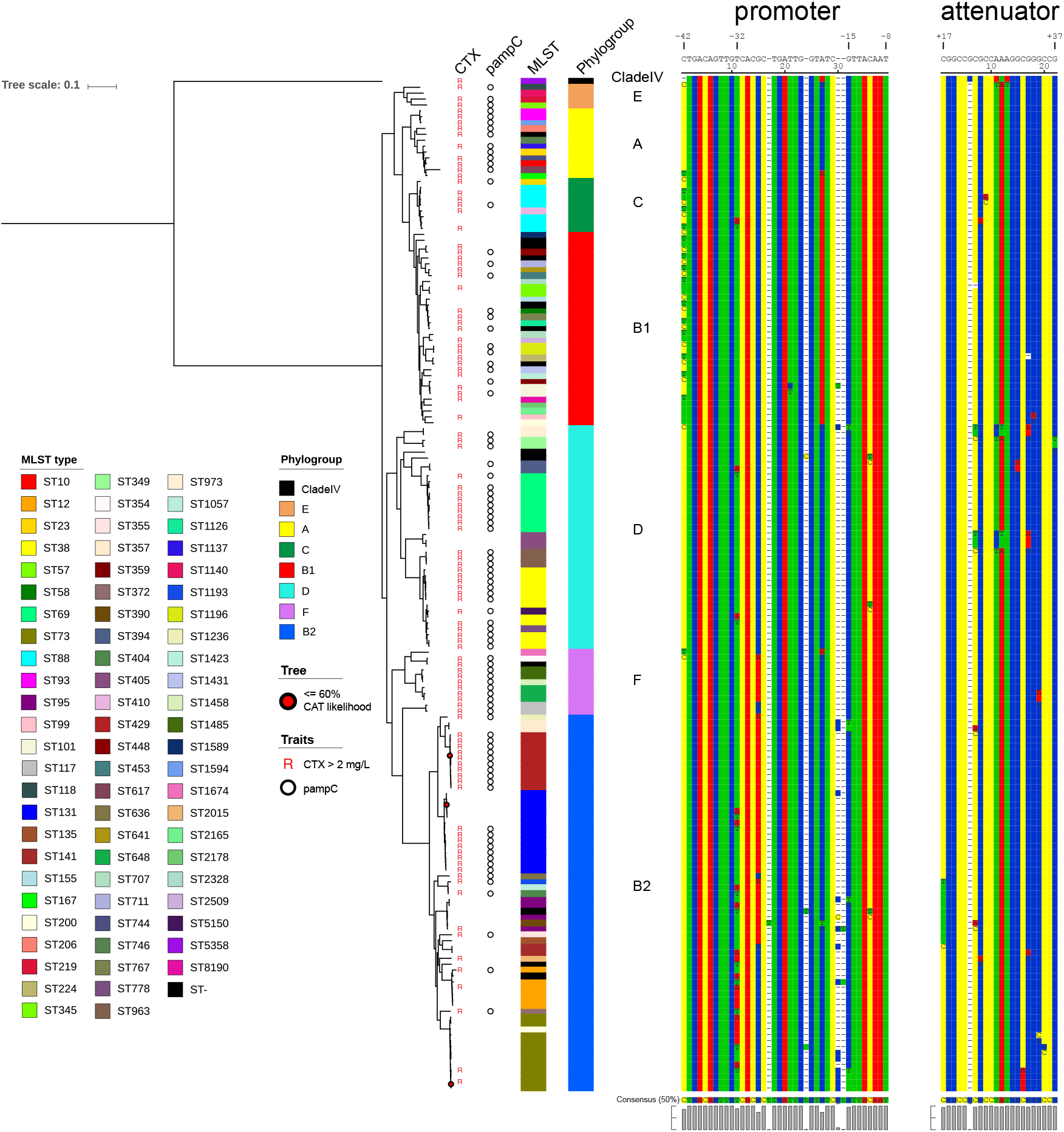
Approximately maximum-likelihood phylogenetic tree of all 172 *E. coli* isolates based on the coreSNP alignment with the resistance for cefotaxime (CTX), *pampC* gene presence, MLSTs, phylogroups, and the alignments of the promoter and the attenuator region. Positions with a CAT likelihood score ≤60% are indicated as red dots.

### Genomic homoplastic mutations

We calculated the Consistency Index for all positions on the *E. coli* reference chromosome. A low Consistency Index value for a position indicates a high degree of inconsistency with the chromosomal phylogeny and can be calculated by HomoplasyFinder as described in earlier studies [10,39,40]. As can be observed in Figure 3, results clearly indicate that notwithstanding multiple other low scoring Consistency Index positions in the promoter and attenuator, position −42 (4,470,140) and −32 (4,470,150) are the lowest scoring, respectively 0.05882353 and 0.07142857 (see also Fig. S1 in the supplemental material). To access how extreme these Consistency Index values are, we calculated the Consistency Index for all positions in the chromosome (see Fig. S2 in the supplemental material). All Consistency Indexes < 1.0 are plotted in the outer ring (ring A) of Figure 4. Results show that only 9,640 out of 5,056,572 positions (0.19%) had a Consistency Index ≤ 0.07142857 (see Figure 4 ring A, cutoff is indicated by black circle). This clearly indicates that positions with low Consistency Indexes are rare, but not unique. Although these 9,640 positions have a low Consistency Index, we do not yet know their relation to CTX resistance.

**FIG 3.**
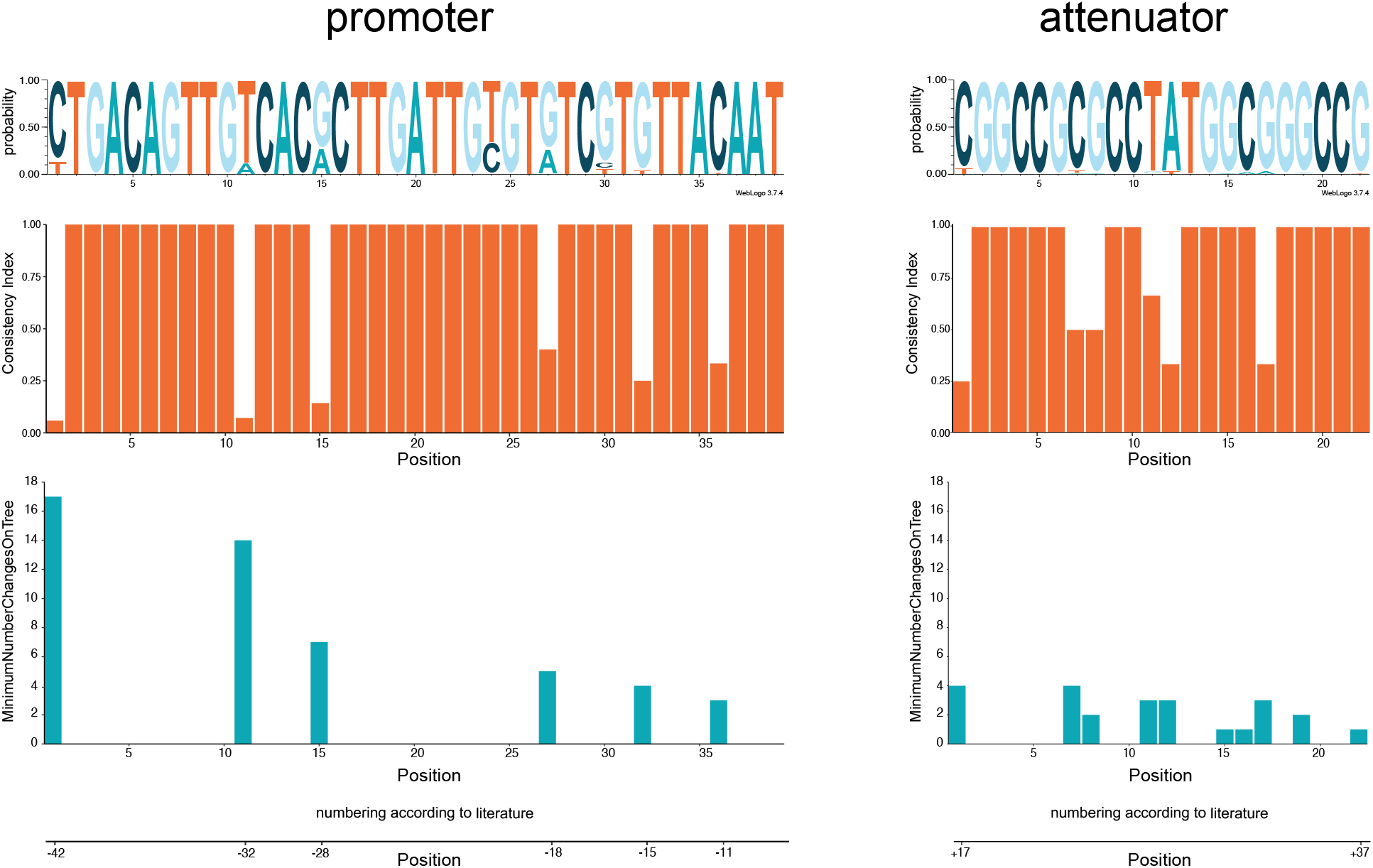
Sequence logo with probability score for the promoter and the attenuator region. The Consistency Index and the minimum number of changes on the tree per position are represented below the sequence logos.

**FIG 4.**
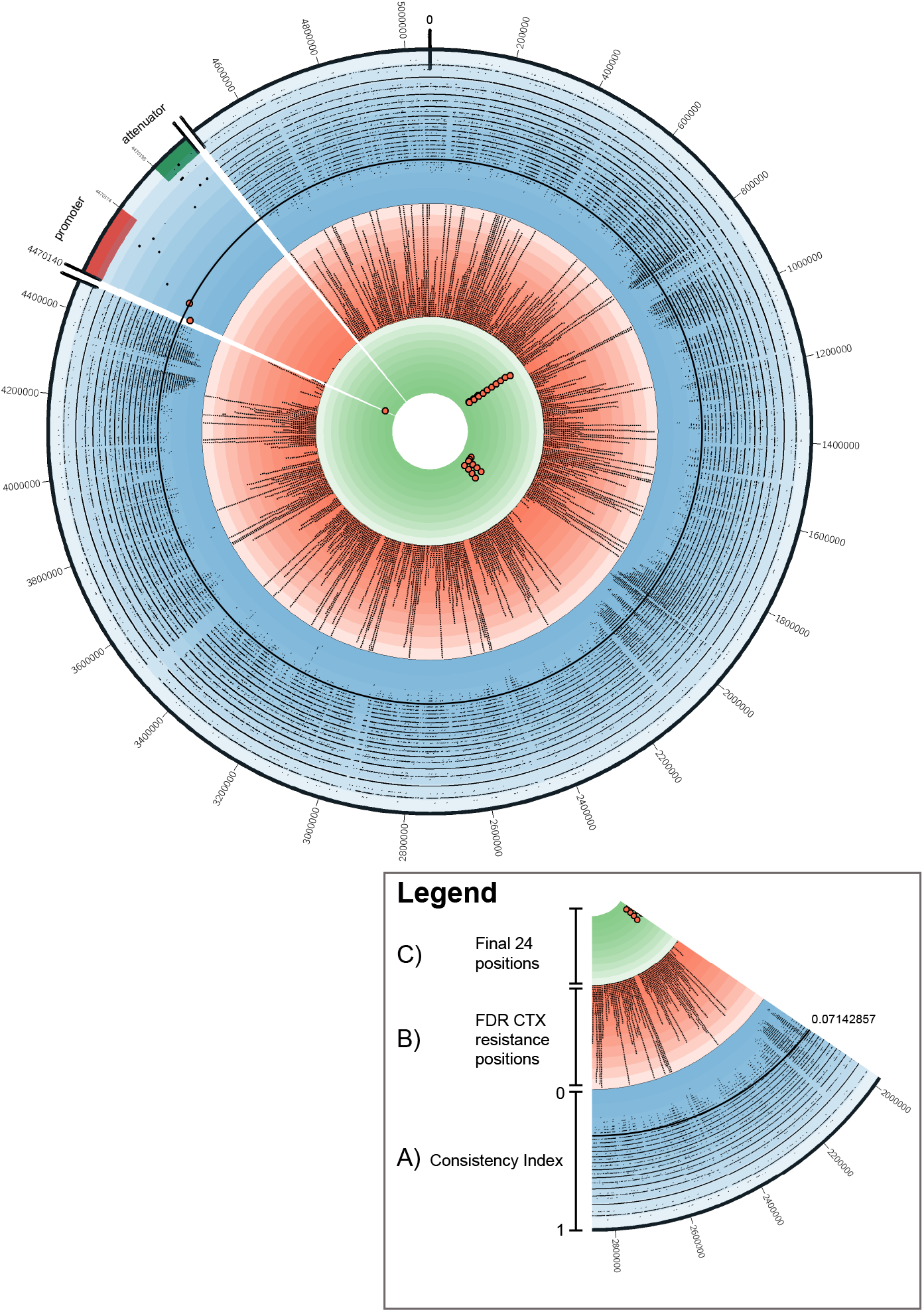
Circos plot for the full chromosome of ampC_0069 (accession no. CP046396) with per position the various metrics used. A) The blue colored ring represents the Consistency Index results per genomic position. The two red dots indicate the −42 and −32 position on the promoter. The black circle line indicates the 0.07142857 Consistency Index value. B) The ring with a red background shows all positions that were significantly associated to cefotaxime (CTX) resistance in all non-*pampC* harboring *E. coli* isolates. Larger bars pointing outwards indicate multiple significant associated positions in a small genomic region. C) The ring with the green background shows all 24 positions that have a low Consistency Index of ≤ 0.05882353 and are significantly associated with CTX resistance in all *non-pampC* harboring *E. coli* isolates.

### CTX resistance measurements

Cefotaxime MIC measurements from Coolen *et al*. in relation to the genotype of the *E. coli* isolates are shown in supplementary table 1. Eighty-four of ninety (93.3%) *pampC* harboring *E. coli* were CTX resistant (MIC >2 mg/L) based on EUCAST clinical breakpoints. Twenty-one of fifty-nine (35.6%) isolates categorized as hyperproducers based on Trazc *et al*. were CTX resistant, primarily isolates with the −42 (*n*=15) or −32 mutation (*n*=2). The *pampC* genes never occurred simultaneously with the −42 or −32 mutations in any of these isolates. One of twenty-three (4.3%) isolates categorized as a low-level AmpC producer (no *pampC* gene and no known ampC promoter mutation) tested CTX resistant and contained an insertion in the spacer region (−16_-15insT), which has not been described by Tracz *et al* [6]. As depicted in Figure 2 the non-*pampC* strains with a phenotype of CTX > 2 mg/L were present in all phylogroups, although CTX resistant isolates with the −42 or −32 mutation were predominantly present in phylogroups B1, B2, and C.

### Geno- to phenotype

To be able to link *E. coli* chromosomal mutations to CTX resistance we excluded all *E. coli* isolates with a plasmid containing an *ampC* β-lactamase gene. The association of SNPs to CTX resistance phenotype (MIC > 2 mg/L) was tested in the remaining 82 isolates using Fisher’s Exact Test. After FDR correction to 0.05, 45,998 significant positions were found (see Figure 4 ring B). Mutation C>T on position −42 of the *ampC* promoter was found to be significantly associated to CTX resistance (FDR = 0.034). However, position −32 A>T was not significantly associated to CTX resistance (FDR = 1).

### Homoplasy-based association analysis

Combining the outcome of the homoplasy analysis with the significant CTX resistance associated positions results in genomic positions associated to CTX resistance that have evolved multiple times independently. After selecting the lowest scoring Consistency Index positions, ≤ 0.05882353, 24 relevant genomic positions were identified that had both a low Consistency Index and a significant association with CTX resistance. Most notably, one of these 24 positions is position −42. Only two mutations of those 24 that were located in genes were non-synonymous: a (conservative) missense mutation in the type II secretion system protein L (*gspL*) gene leading to Ser330Thr alteration and a mutation in the hydroxyethylthiazole kinase (*thiM*) gene resulting in a Thr122Ala alteration according to the annotation of *E. coli* strain ampC_0069 (accession no. CP046396). In addition to the non-synonymous mutation found on the *gspL* gene, eight synonymous mutations are also located in genes annotated as being part of the type II secretion system. A complete overview is presented in Table 2.

**TABLE 2.**
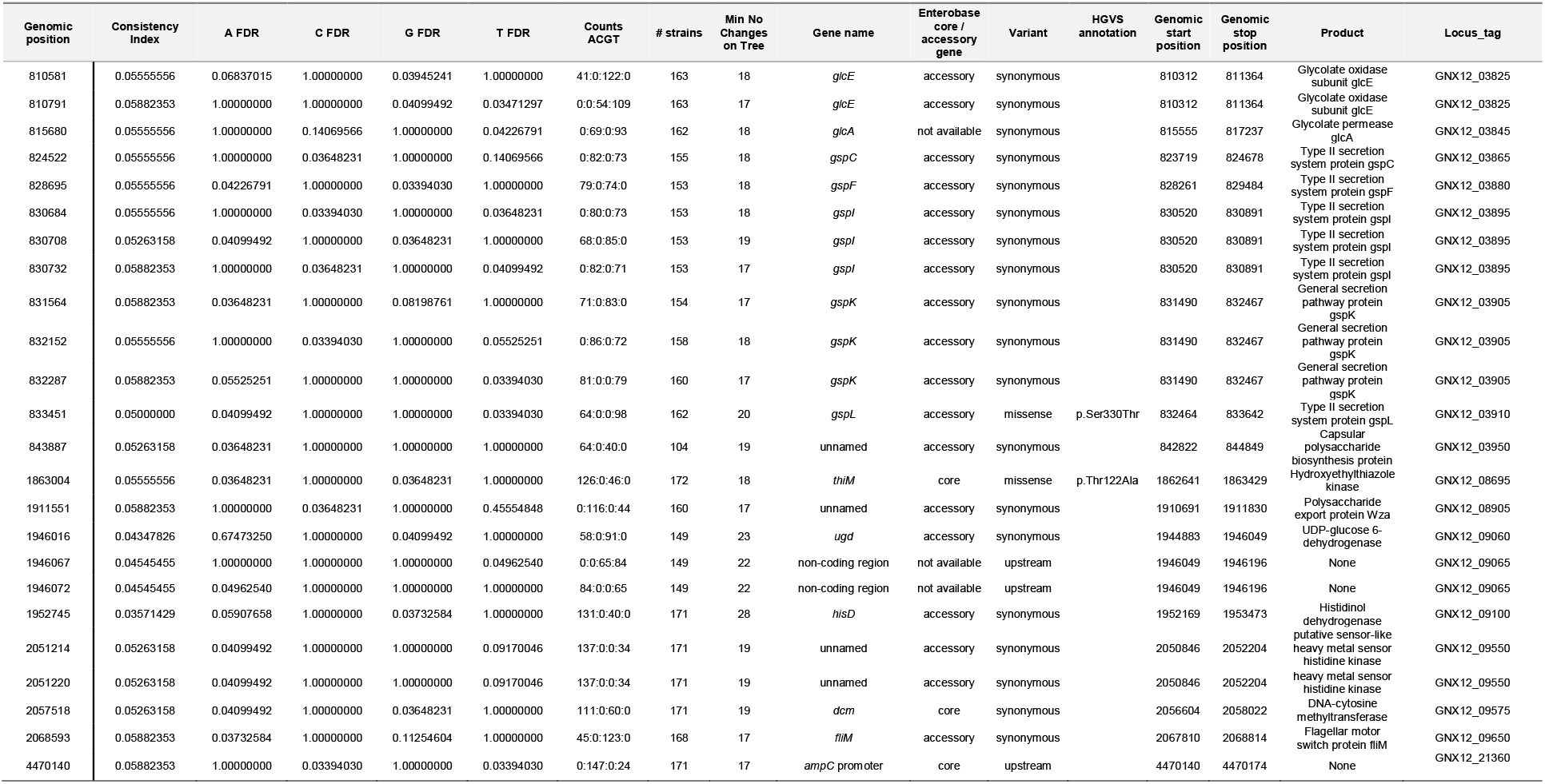
The *n=24* positions with a significant association with cefotaxime resistance (FDR ≤0.05) and with a consistency index ≤ 0.05882353.

### Recombination analysis

To verify if the level of homoplasy could be a result of recombination, we used Genealogies Unbiased By recomBinations In Nucleotide Sequences (Gubbins) algorithm to predict recombination events in our isolate collection [34]. This analysis showed frequent recombination events in our 172 *E. coli* isolates (see Fig. S3 in the supplemental material). Results illustrate that recombination blocks cover the region of the *gspL* and the *thiM* gene and their high homoplasy levels could thus be due to recurrent recombination rather than independent mutations. Nonetheless, position −42 in the *ampC* promoter is not located in a region effected by recombination as shown in Fig. S3. Moreover, when inferring the phylogenetic tree corrected for recombination events as obtained from the Gubbins analysis, the −42C>T mutation actually occurred in 18 independent branches rather than the 17 branches in the uncorrected tree. This supports our previous results that this mutation is homoplastic, and not the results of a recurrent recombination event.

## Discussion

We present a genome-wide analysis in which homoplastic mutations are associated with antibiotic resistance in *E. coli*. By comparing whole-genome sequencing data of 172 *E. coli* isolates to a tailored reference chromosome we were able to reconstruct the evolution of the genomes and therewith map recurrent events, allowing us to detect homoplasy associated to CTX resistance.

Our foremost finding is the significant association of the −42C>T mutation, in the *ampC* promoter, to CTX resistance that evolved independently at least 17 times in 5 distinct phylogroups. The −42C>T mutation has been confirmed in former studies to result in AmpC hyperproduction in *E. coli*. Nelson *et al*. demonstrated an 8 to 18 times increase in activity of AmpC when cloning the promoter upstream a *lac* operon [41]. Vice versa, Caroff *et al*. found a decrease in expression of AmpC when cloning the promoter with a −42T>C mutation in a pKK232-8 reporter plasmid with chloramphenicol acetyltransferase gene [24]. Tracz *et al*. confirmed that the −42C>T mutation has the strongest effect on the *ampC* promoter, resulting in a high expression of the *ampC* gene as detected by RT-qPCR [6]. Despite the fact that the −42C>T mutation has such a strong effect on AmpC production the effect of the mutation on CTX MICs had not been confirmed. Moreover, the contribution of convergent evolution on this position relative to the role of the expansion of a clone with a beneficial mutation at this position has not been determined. That being the case, this study provides evidence that this −42 C>T mutation is not a result of a recombination event and most likely evolved many times independently. Remarkably, we observed that the −42C>T mutation never occurs in the presence of a *pampC* gene (in zero out of twenty-four cases). This was even noticed in isolates with the same MLST, i.e. ST88 −42C>T (*n*=3) and *pampC* (*n*=1), suggesting preferred exclusivity for one of the resistance mechanisms. One study mentioned the co-occurrence of the −42C>T mutation and a *pampC* gene in only one out of thirty-six strains [42]. One could speculate that the exclusivity is a matter of what arrives first, the plasmid or the mutation, after which there is no selective advantage for the second mechanism, or that there is actually a fitness cost to having both the mutation and the plasmid relative to having only the mutation or the plasmid.

The study performed by Tracz *et al*. showed that position −32T>A on the promotor of *ampC* associates with AmpC hyperproduction that results in elevated MIC levels for FOX [6]. Surprisingly, in the current study no significant association of −32T>A with CTX resistance was noticed despite its low Consistency Index. Only two out of twenty isolates with the −32T>A were CTX resistant, four out of twenty showed an intermediate elevated CTX MIC, and fourteen were susceptible for CTX. Although we do not know under which conditions this mutation did arise, it can be speculated that the high level of homoplasy at the −32 position is associated with a different trait, e.g. resistance against another antibiotic.

Prior studies discovered the importance of mutations in the promoter elements. Random sequences can even evolve expression comparable to the wild-type promoter elements after only a single mutation [43]. Furthermore, these promotor elements evolve to only a few forms indicating convergent evolution [44], as also observed in the present study. All encountered variants seem to result in a sequence that resembles the *E. coli* consensus sigma 70 promoter more than the wild type sequence they are derived from [6].

Next to the −42C>T promoter mutation we detected twenty-three other positions in our analysis that are associated with CTX resistance and have extreme high levels of homoplasy. Most of these are synonymous mutations, with only two missense mutations (*thiM* and *gspL*) found. It is remarkable that one missense mutation (p.Ser330Thr) is located in *gspL* that encodes for a protein of the type II secretion system. The type II secretion system is used by many gram-negative bacteria to translocate folded proteins from the periplasm, through the outer membrane, into the extracellular milieu [45]. The system is composed of 12–15 different general secretory pathway (Gsp) proteins and is related to virulence of various pathogenic *E. coli*, e.g. EHEC and UPEC [46–48]. It could be that in our selection of mainly clinical samples a certain predilection has occurred towards isolates with particular virulence traits. The *gspL* gene has been described as being part of the accessory genome of *E. coli* [49]. Our study supports this finding as some strains did not harbor this gene. Additionally, we found evidence that recombination events in the type II secretion system could be the underlying cause of the extreme homoplasy levels. Still, it is remarkable that missense mutation p.Ser330Thr in the *gspL* gene correlates with the CTX resistance trait even though it is most likely caused by a recombination event. To the best of our knowledge no relationship between type II secretion system and CTX resistance has been observed before. One could hypothesize that the mutation is a secondary adaptation needed to cope with the elevated AmpC production, as the peptidoglycan (PG) layer is effected by AmpC hyperproduction and the type II secretion system contains proteins that are partly localized in the periplasm [50,51].

The use of genomic data to detect homoplasy events is not an uncommon scientific technique [52–54]. In *Mycobacterium tuberculosis*, it is a well-known method to identify advantageous mutations, as they are likely to be associated with phenotypes such as drug resistance, heightened transmissibility, or host adaptation [12–15]. A similar approach was taken recently by Benjak *et al*. to screen for highly polymorphic genes and genomic regions of *Mycobacterium leprae* [55]. Homoplasy-based association analysis limits phylogenetic bias by correcting for genetic relatedness of strains with the same phenotype, thereby increasing statistical power to find true associations [14]. Taking this into account, the use of homoplasy-based association analysis seems viable to relate polymorphic sites to phenotypic traits in bacteria. Still, studies on other genera than mycobacteria are scarce. To our knowledge, no homoplasy studies have used this method on *E. coli*.

The increase of 3GC resistance imposes a clinical threat by restricting treatment options and it is essential to understand the underlying resistance mechanisms. To be able to explore these mechanisms we selected primarily clinical *E. coli* strains. The current study is directed on exploring AmpC mediated CTX resistance. Therefore, we included isolates that are already suspected for increased AmpC production based on elevated FOX resistance. Since a random sample of *E. coli* would limit finding homoplasy-based associated promoter mutations with CTX resistance. A downside of these selection criteria might be that we over-estimated certain genetic variants associated with the trait, as we do not know the frequency of these variants in the general population. Despite the fact that the spontaneous mutation rate in *E. coli* is relatively low [56], it is still likely that this particular mutation occurs often in the general population, given the vast amounts of *E. coli* in the environment [57], providing ample opportunities for adaptation to antibiotics and arguing for antibiotics of which genomic adaptation requires multiple mutations in order to develop resistance.

Findings of this study have a number of implications for future practice. This study not only grants insights into how chromosomal-encoded antibiotic resistance evolves, but also provides potential strategies for future homoplasy-based association studies. Furthermore, the use of genome-wide homoplasy-based analysis could be applied to optimize outbreak analysis. Prior studies have optimized outbreak analysis by removing recombinant regions [58, 59]. Homoplasy events disturbs the true phylogeny, hence, removing genomic positions which are heavily affected by homoplasy could improve tree topology, thereby refining outbreak analysis, although this strategy is still under debate [60].

## Conclusions

To conclude, our method demonstrates extreme levels of homoplasy in *E. coli* that are significantly associated with CTX resistance. Greater access to WGS data provides new opportunities to perform large-scale genome-wide analysis. Homoplasy-based methods can have a potential role in future studies as they constitute an effective strategy to relate phenotypic traits to variable genomic positions.

## Abbreviations

3GC: third-generation cephalosporin
*campC*: chromosomal-mediated ampC
CAT: computerized adaptive testing
CTX: cefotaxime
DNA: Deoxyribonucleic acid
EHEC: enterohemorrhagic *Escherichia Coli*
ESBL: extended-spectrum β-lactamases
FOX: cefoxitin
gDNA: genomic DNA
MICs: minimal inhibitory concentrations
MLST: multilocus sequence typing
*pampC*: plasmid-mediated ampC
PG: peptidoglycan
qRT-PCR: quantitative reverse transciptase polymerase chain reaction
SMRT: Single-molecule real-time sequencing
SNP: single-nucleotide polymorphism
ST: sequence type
UPEC: uropathogene *Escherichia Coli*
WGS: whole genome sequencing

## Author contributions

HW, JK, and MH conceived and supervised the study. JC, ED, EK, JS, JV, WM, and KN performed the data acquisition. JC and ED performed the data analysis. JC performed bioinformatic analysis. JC, ED, and MH performed the data interpretation and wrote the manuscript. All authors read and approved the final manuscript. All authors read and approved the final manuscript.

## Acknowledgements

Special thanks to A. C. J. Soer (Department of Medical Microbiology and Radboudumc Center for Infectious Diseases, Radboudumc, Nijmegen, the Netherlands), B. A. Lamberts (Department of Medical Microbiology and Immunology, Rijnstate, Arnhem, the Netherlands) and C. Verhulst (Department of Infection Control, Amphia Ziekenhuis, Breda, The Netherlands and Laboratory for Microbiology, Microvida, Location Breda, The Netherlands) for handling the samples on the lab and creating the Illumina sequence libraries.

M. P. Kwint and R. Derks (Department of Human Genetics, Radboudumc, Nijmegen, the Netherlands) for helping with SMRT sequencing on the PacBio Sequel I.

Many thanks to M. Janssens (Laboratory for Medical Microbiology and Immunology, Elisabeth-Tweesteden Hospital, Tilburg, the Netherlands), S. Van Leest (Laboratory for Microbiology, Microvida, location Bravis Hospital, the Netherlands), K. T. Veldman and D. J. Mevius (department of Bacteriology and Epidemiology, Wageningen Bioveterinary Research, Lelystad, the Netherlands, E.A. Reuland (Medical Microbiology and Infection Control, Amsterdam UMC location VUmc, Amsterdam, the Netherlands), W. H. F. Goessens (Erasmus University Medical Center, Rotterdam, Netherlands), R. W. Bosboom (Department of Medical Microbiology and Immunology, Rijnstate, Arnhem, the Netherlands) and P. Vos (Check-Points, Wageningen, the Netherlands) for providing samples of which some are included in this study.

---

**FIG S1** Violin plots of the log10 Consistency Indexes of the promoter and attenuator.

**FIG S2** Distribution of the log10 Consistency Indexes of all genomic position based on the *E. coli* ampC_0069 reference chromosome, compared to the log10 Consistency Indexes of the promoter and attenuator region.

**FIG S3** Recombination events inferred from all 172 *E. coli* isolates by Gubbins displayed along the approximately maximum-likelihood phylogenetic tree based on the coreSNP alignment. Phylogroups are depicted as in FIG 2. Gubbins blocks are colored red if they are ancestral, and blue if they only affect one isolate. The line graph represents the recombination prevalence along the sequence. The 24 positions with a significant association with cefotaxime resistance (FDR ≤0.05) and a consistency index ≤ 0.05882353 are indicated on the top of the figure. The two missense mutations and *ampC* promoter region are displayed in blue.

**Table S1** Classification of n=172 *E. coli* isolates in the three genotypes (*pampC*, hyperproducer, low-level AmpC producers), with the results of the MLST and phylogroups stratification and the different mutations in the promoter and attenuator per isolate. Isolates with a CTX MIC >2 mg/L without a confirmed *pampC* gene are depicted in **bold**.

**Table S2** SNP analysis for n=172 *E. coli* isolates according to snippy statistics.

