## Supplementary figures and images for "Genome-wide analysis in *Escherichia coli* unravels an unprecedented level of genetic homoplasy associated with cefotaxime resistance"

### Supplemental Figure 1

**FIG S1** Violin plots of the log10 Consistency Indexes of the promoter and attenuator.

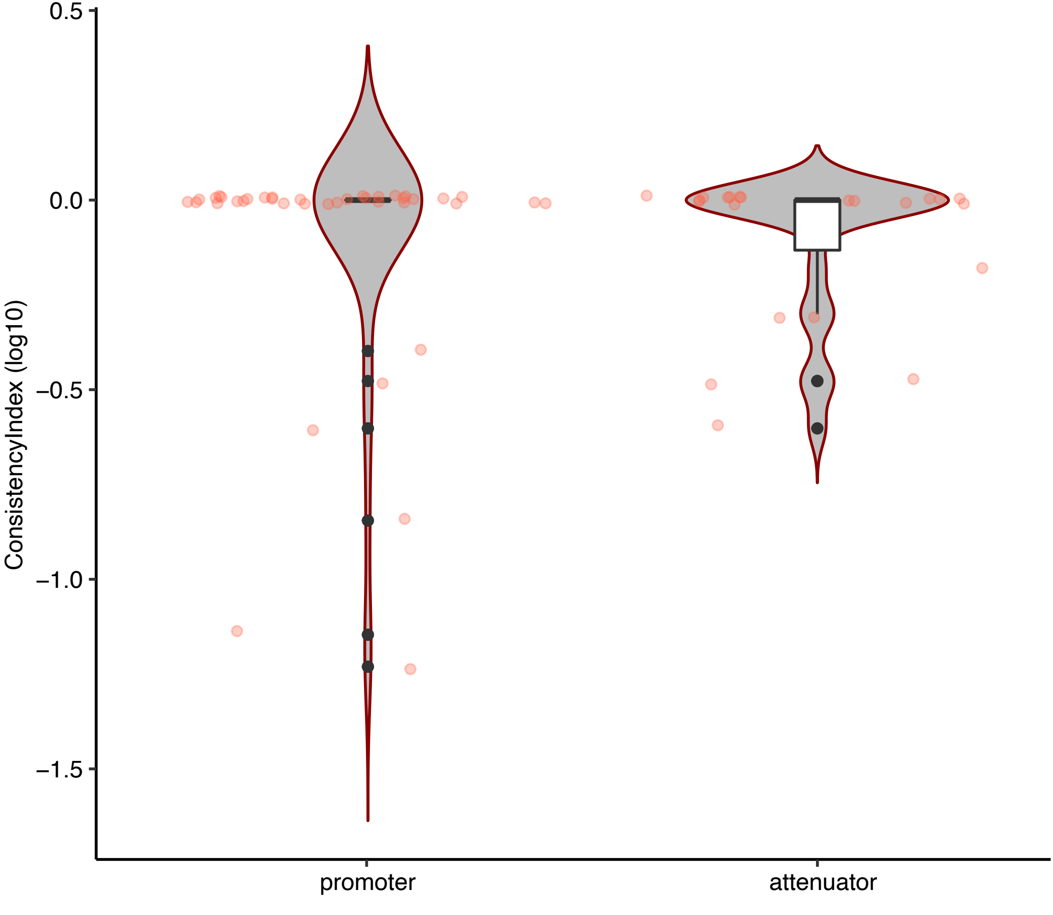
