## Supplemental Figure 2 for "Genome-wide analysis in *Escherichia coli* unravels an unprecedented level of genetic homoplasy associated with cefotaxime resistance"

**FIG S2** Distribution of the log10 Consistency Indexes of all genomic position based on the *E. coli* ampC_0069 reference chromosome, compared to the log10 Consistency Indexes of the promoter and attenuator region.

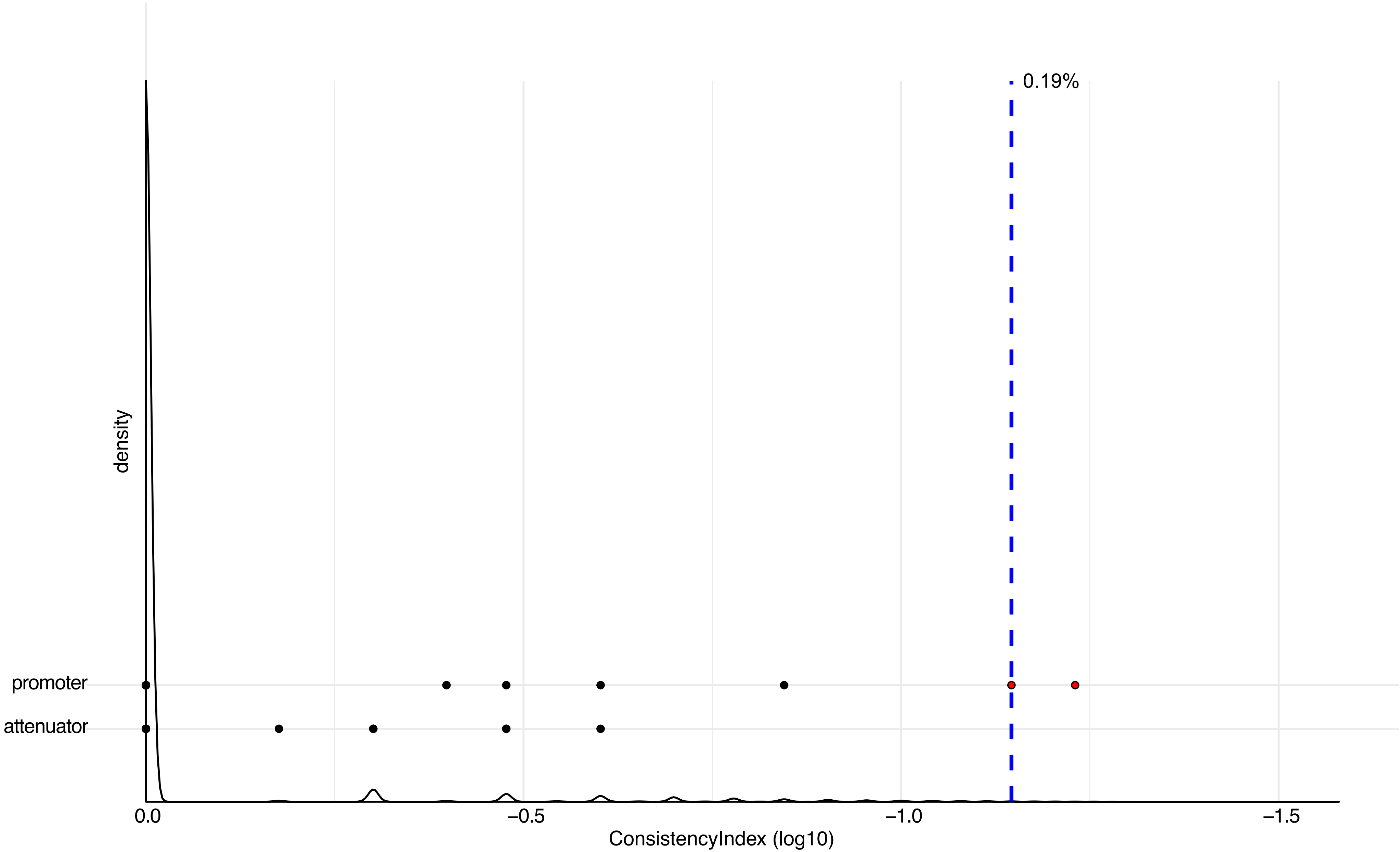
