## Supplemental Figure 3 for "Genome-wide analysis in *Escherichia coli* unravels an unprecedented level of genetic homoplasy associated with cefotaxime resistance"

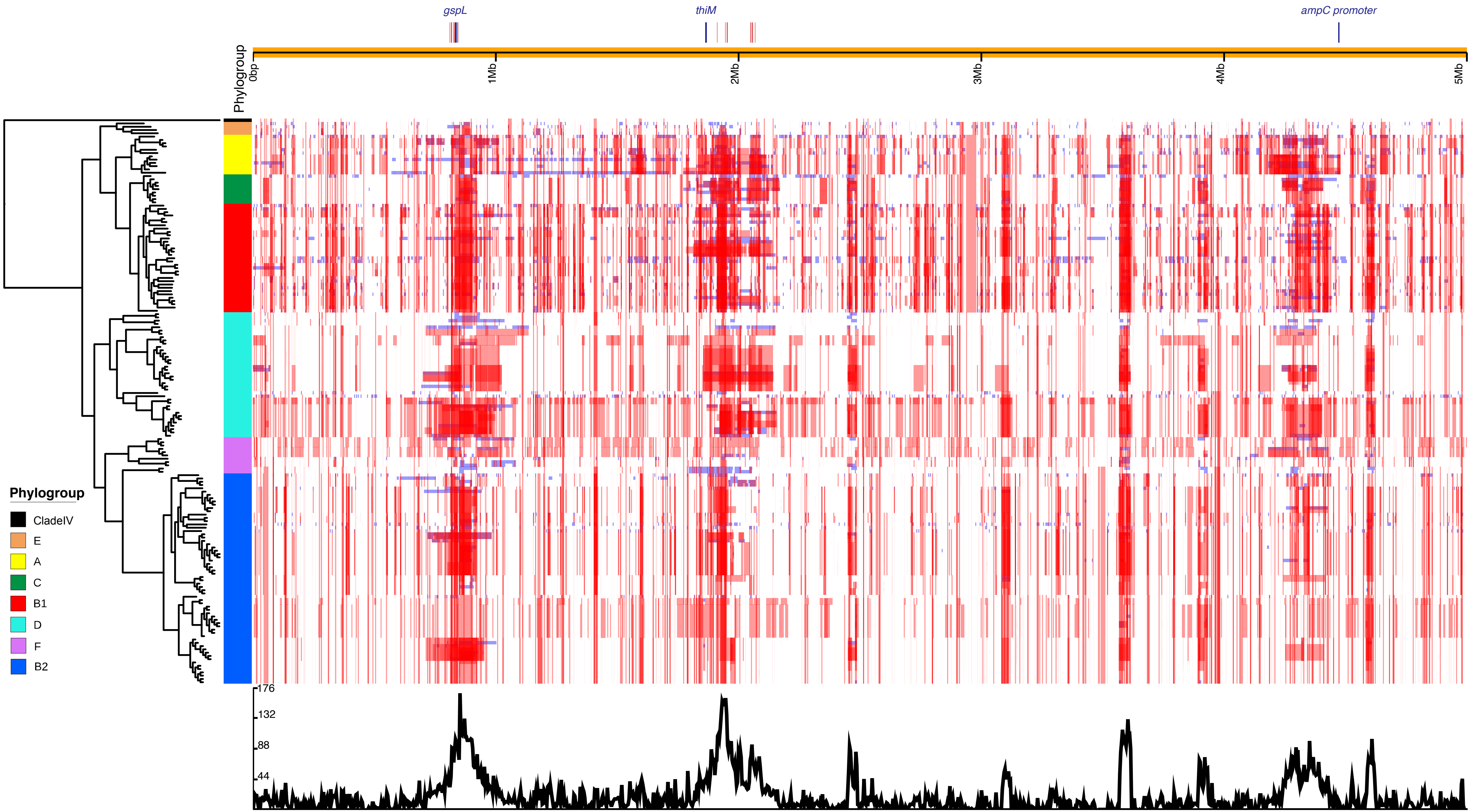


**FIG S3** Recombination events inferred from all 172 *E. coli* isolates by Gubbins displayed along the approximately maximum-likelihood phylogenetic tree based on the coreSNP alignment. Phylogroups are depicted as in FIG 2. Gubbins blocks are coloured red if they are ancestral, and blue if they only affect one isolate. The line graph represents the recombination prevalence along the sequence. The 24 positions with a significant association with cefotaxime resistance (FDR ≤0.05) and a consistency index ≤ 0.05882353 are indicated on the top of the figure. The two missense mutations and *ampC* promoter region are displayed in blue.
