## Supplemental Table 1 for "Genome-wide analysis in *Escherichia coli* unravels an unprecedented level of genetic homoplasy associated with cefotaxime resistance"

**Table S1**

Classification of $n$=172 *E. coli* isolates in the three genotypes based on Tracz *et al.*[5] (pampC, hyperproducer, low-level), with MIC for CTX (mg/L), the results of the MLST and phylogroups stratification and the different mutations in the promoter and attenuator per isolate. Isolates with a CTX MIC >2 mg/L without a confirmed *pampC* gene are depicted in **bold**.

| | | | | | Promoter | | | | | | | | | | | | Attenuator | | | | | | | | | | | | | |
|---|---|---|---|---|---|---|---|---|---|---|---|---|---|---|---|---|---|---|---|---|---|---|---|---|---|---|---|---|---|---|
| | | | | | displaced -35 box | | -35 box | | | | | displaced 10 box | Spacer | | | -10 box | | | | | | | | | | | | | | |
| | | | | | -44 | -42 | -32 | -28 | -25/-26 | -23 | -21/-20 | -18 | -16/-15 | -15/-14 | -13/-12 | -11 | +17 | +21/+22 | +22 | +23 | +24 | +26 | +27 | +28 | +30 | +32 | +34 | +35 | +36 | +37 |
| unique_ID | ampC genotype | MIC CTX (mg/L) | MLST | Phylogroup | T | C | T | G | - | G | - | G | - | - | - | C | C | - | C | G | C | T | A | T | G | G | G | C | C | G |
| ampC_0001 | hyperproducer | 0.38 | 357 | B2 | | | | A | | | | | | T | | | | | | | | | | | | | | | | |
| ampC_0002 | low-level | 0.38 | 38 | D | | | | | | | | | | | | T | | | | | | | | | | | | | | |
| ampC_0003 | low-level | 0.38 | 405 | D | | | | | | | | | | | | | | | T | | | G | T | | | A | | | | |
| ampC_0004 | hyperproducer | 1.5 | 73 | B2 | | | A | | | | | | | | | | | | | | | | | | | | | | | |
| ampC_0005 | hyperproducer | 0.5 | 131 | B2 | | | | A | | | | | | G | | | | | | | | | | | | | | | | |
| ampC_0006 | low-level | 0.25 | 405 | D | | | | | | | | | | | | | | | T | | | G | T | | | A | | | | |
| ampC_0007 | hyperproducer | 0.5 | 73 | B2 | | | A | | | | | | | | | | | | | | | | | | | | | | | |
| **ampC_0008** | **hyperproducer** | **3** | **1126** | **B1** | | T | | | | | | A | | | | | | | | | | | | | | | | | | |
| ampC_0009 | blaCMY-2 | 12 | 69 | D | | | | | | | | | | | | | | | | | | | | | | | | | | |
| ampC_0010 | blaCMY-2 | 4 | 1431 | B1 | | | | | | | | A | | | | | | | | | | | | | | | | | | |
| ampC_0011 | blaCMY-2 | 12 | 131 | B2 | | | | A | | | | | | | | | | | | | | | | | | | | | | |
| ampC_0012 | hyperproducer | 1 | 12 | B2 | | | A | | | | | | | | | | | | | | | | | | | | | | | |
| ampC_0013 | blaDHA-1 | 2 | 69 | D | | | | | | | | | | | | | | | | | | | | | | | | | | |
| ampC_0014 | blaDHA-1 | 3 | 38 | D | | | | | | | | | | | | | | | | | | | | | | | | | | |
| ampC_0015 | low-level | 0.25 | 746 | A | | | | | | | | | | | | | | | | | | | | | | | | | | |
| ampC_0016 | low-level | 0.19 | 1589 | B1 | | | | | | | | A | | | | | | | | | | | | | | | | | | |
| ampC_0017 | hyperproducer | 0.5 | 12 | B2 | | | A | | | | | | | | | | | | | | | | | | | | | | | |
| ampC_0018 | low-level | 0.19 | 131 | B2 | | | | A | | | | | | | | | | | | | | | | | | | | | | |
| ampC_0019 | low-level | 0.094 | 155 | B1 | | | | | | | | A | | | | | | | | | | | | | | | | | | |
| ampC_0020 | low-level | 0.125 | 1140 | E | | | | | | | | | | | | | | | | | | | | | | | | | | |
| ampC_0021 | blaCMY-2 | 6 | 636 | B2 | | | | | | | | | | | | | | | | | | | | | | | | | | |
| ampC_0022 | blaCMY-2 | 32 | 1485 | F | | | | A | | | | | | | | | | | | | | | | | | | | | | |
| ampC_0023 | blaCMY-42 | 32 | 617 | A | | | | | | | | | | | | | | | | | | | | | | | | | | |
| ampC_0024 | low-level | 0.38 | 88 | C | | | | | | | | A | | | | | | | | | | | | | | | | | | |
| ampC_0025 | low-level | 0.38 | 131 | B2 | | | | A | | | | | | | | | | | | | | | | | | | | | | |
| ampC_0026 | blaCMY-2 | 16 | 354 | F | | | | A | | | | | | | | | | | | | | | | | | | | | | |
| ampC_0027 | blaCMY-2 | 32 | 778 | D | | | | | | | | | | | | | | | | | | | | | | | | | | |
| ampC_0028 | hyperproducer | 0.5 | 73 | B2 | | | | | | | | | | T | | | | | | | | | | | | | | | | |
| ampC_0029 | blaCMY-2 | 32 | 69 | D | | | | | | | | | | | | | | | | | | | | | | | | | | |
| ampC_0030 | blaCMY-2 | 24 | 1458 | F | | | | A | | | | | | | | | | | | | | | | | | | | | | |
| ampC_0031 | blaCMY-2 | 4 | 10 | A | | | | | | | | | | | | | | | | | | | | | | | | | | |
| ampC_0032 | low-level | 0.19 | 131 | B2 | | | | A | | | | | | | | | | | | | | | | | | | | | | |
| ampC_0033 | low-level | 0.19 | 405 | D | | | | | | | | | | | | | | | T | | | G | T | | | A | | | | |
| ampC_0034 | low-level | 0.38 | unknown ST | D | | | | | | | | | | | | | | | | | | | | | | | | | | |
| ampC_0035 | low-level | 0.38 | 88 | C | | | | | | | | A | | | | | | | | | | | | | | | | | | |
| **ampC_0036** | **hyperproducer** | **4** | **224** | **B1** | | T | | | | | | A | | | | | | | | | | | | | | del | | | | |
| ampC_0037 | blaCMY-2 | 12 | unknown ST | B1 | | | | | | | | A | | | | | | | | | | | | | | | | | | |
| ampC_0038 | blaDHA-1 | 2 | 372 | B2 | | | | | | | | | | | | | | | | | | | | | | | | | | |
| ampC_0039 | blaCMY-2 | 16 | 744 | A | | | | | | | | | | | | | | | | | | | | | | | | | | |
| ampC_0040 | blaCMY-2 | 12 | 429 | B2 | | | | A | | | | | | | | | | | | | | | | | | | | | | |
| ampC_0041 | blaCMY-2 | 32 | 23 | C | | | | | | | | A | | | | | | | | | | | | | | | | | | |
| ampC_0042 | hyperproducer | 2 | 73 | B2 | | | | | | | | | | T | | | | | | | | | | | | | | | | |
| ampC_0043 | low-level | 0.125 | unknown ST | B2 | | | | | | | | | | | | | | | | | | | | | | | | | | |
| ampC_0044 | blaCMY-2 | 32 | 648 | F | | | | A | | | | | | | | | | | | | | | | | | | | | | |
| ampC_0045 | blaCMY-2 | 24 | 131 | B2 | | | | A | | | | | | | | | | | | | | | | | | | | | | |
| ampC_0046 | hyperproducer | 1.5 | 2178 | B1 | | T | | | | | | A | | | | | | | | | | | | | | | | | | |
| **ampC_0047** | **hyperproducer** | **3** | **2509** | **B1** | | T | | | | | | A | | | | | | | | | | | | | | | | | | |
| ampC_0048 | hyperproducer | 0.75 | 345 | B1 | | T | | | | | | A | | | | | | | | | | | | | | | | | | |
| ampC_0049 | blaCMY-2 | 32 | 219 | E | | | | | | | | | | | | | | | | | | | | | | | | | | |
| ampC_0050 | blaCMY-2 | 32 | 963 | D | | | | | | | | | | | | | | | | | | | | | | | | | | |
| **ampC_0051** | **hyperproducer** | **3** | **410** | **C** | | T | | | | | | A | | | | | | | | | | | | | | | | | | |
| **ampC_0052** | **hyperproducer** | **4** | **88** | **C** | | T | | | | | | A | | | | | | | | | | | | | | | | | | |
| ampC_0053 | low-level | 0.25 | 69 | D | | | | | | | | | | | | | | | | | | | | | | | | | | |
| ampC_0054 | blaCMY-2 | 32 | unknown ST | F | | | | A | | | | | | | | | | | | | | | | | | | | | | |
| ampC_0055 | hyperproducer | 2 | 2165 | B1 | | T | | | | | | A | | | | | | | | | | | | | | | | | | |
| ampC_0056 | blaCMY-2 | 12 | 349 | D | | | | | | | | | | | | | | | | | | | | | | | | | | T |
| ampC_0057 | blaCMY-2 | 16 | 404 | B2 | | | | A | | | | | | | | | T | | | | | | | | | | | | | |
| ampC_0058 | blaCMY-2 | 12 | 69 | D | | | | | | | | | | | | | | | | | | | | | | | | | | |
| ampC_0059 | blaCMY-59 | 12 | 93 | A | | | | | | | | | | | | | | | | | | | | | | | | | | |
| ampC_0060 | hyperproducer | 0.75 | 12 | B2 | | | A | | | | | | | | | | | | | | | | | | | | | | | |
| ampC_0061 | blaCMY-42 | 32 | unknown ST | B1 | | | | | | | | A | | | | | | | | | | | | | | | | | | |
| ampC_0062 | blaCMY-2 | 12 | 429 | B2 | | | | A | | | | | | | | | | | | | | | | | | | | | | |
| ampC_0063 | hyperproducer | 1 | unknown ST | B1 | | T | | | | | | A | | | | | | | | | | | | | | | | | | |
| **ampC_0064** | **hyperproducer** | **3** | **unknown ST** | **B1** | | T | | | | | | A | | | | | | | | | | | | | | | | | | |
| ampC_0065 | hyperproducer | 1 | 73 | B2 | | | A | | | | | | | | | | | | | | | | | | | | C | | | |
| ampC_0066 | blaCMY-2 | 32 | 963 | D | | | | | | | | | | | | | | | | | | | | | | | | | | |
| ampC_0067 | blaCMY-2 | 3 | 1193 | B2 | | | | A | | | | | | | | | T | | | | | | | | | | | | | |
| ampC_0068 | blaCMY-2 | 24 | 1236 | B2 | | | | A | | | | | | | | | | | | | | | | | | | | | | |
| ampC_0069 | blaCMY-42 | 32 | 648 | F | | | | A | | | | | | | | | | | | | | | | | | | A | | | |
| ampC_0070 | blaCMY-2 | 6 | 69 | D | | | | | | | | | | | | | | | | | | | | | | | | | | |
| ampC_0071 | blaCMY-2 | 6 | 429 | B2 | | | | A | | | | | | | | | | | | | | | | | | | | | | |
| ampC_0072 | blaCMY-2 | 12 | 93 | A | | | | | | | | | | | | | | | | | | | | | | | | | | |
| ampC_0073 | blaCMY-2 | 8 | 131 | B2 | | | | A | | | | | | | | | | | | | | | | | | | | | | |
| ampC_0074 | blaCMY-2 | 12 | 69 | D | | | | | | | | | | | | | | | | | | | | | | | | | | |
| ampC_0075 | blaCMY-2 | 1.5 | 359 | B1 | | | | | | | | A | | | | | | | | | | | | | | | | | | |
| ampC_0076 | blaCMY-2 | 12 | 38 | D | | | | | | | | | | | | | | | | | | | | | | | | | | |
| ampC_0077 | blaCMY-2 | 32 | 429 | B2 | | | | A | | | | | | | | | | | | | | | | | | | | | | |
| ampC_0078 | blaCMY-2 | 6 | 1594 | A | | | | | | | | | | | | | | | | | | | | | | | | | | |
| ampC_0079 | blaCMY-2 | 6 | 1137 | A | | | | | | | | | | | | | | | | | | | | | | | | | | |
| ampC_0080 | blaCMY-2 | 8 | 117 | F | | | | | | | | | | | | | | | | | | | | | | | | | | |
| ampC_0081 | blaCMY-2 | 32 | 38 | D | | | | | | | | | | | | | | | | | | | | | | | | | | |
| ampC_0082 | blaCMY-2 | 32 | 448 | B1 | | | | | | | | A | | | | | | | | | | | | | | | | | | |
| ampC_0083 | blaCMY-2 | 32 | 131 | B2 | | | | A | | | | | | | | | | | | | | | | | | | | | | |
| ampC_0084 | blaCMY-2 | 32 | 131 | B2 | | | | A | | | | | | | | | | | | | | | | | | | | | | |
| ampC_0085 | blaCMY-2 | 16 | 131 | B2 | | | | A | | | | | | | | | | | | | | | | | | | | | | |
| ampC_0086 | blaDHA-1 | 2 | 69 | D | | | | | | | | | | | | | | | | | | | | | | | | | | |
| ampC_0087 | blaCMY-2 | 32 | 963 | D | | | | | | | | | | | | | | | | | | | | | | | | | | |
| ampC_0088 | blaCMY-2 | 12 | 38 | D | | | | | | | | | | | | | | | | | | | | | | | | | | |
| ampC_0089 | blaCMY-2 | 8 | 206 | A | | | | | | | | | | | | | | | | | | | | | | | | | | |
| ampC_0090 | blaCMY-2 | 4 | 58 | B1 | | | | | | | | A | | | | | | | | | | | | | | | | | | |
| ampC_0091 | blaCMY-2 | 4 | 767 | B1 | | | | | | | | A | | | | | | | | | | | | | | | | | | |
| ampC_0092 | blaCMY-2 | 8 | 57 | E | | | | | | | | | | | | | | | | | | | | | | | | | | |
| ampC_0093 | blaCMY-2 | 8 | 429 | B2 | | | | A | | | | | | | | | | | | | | | | | | | | | | |
| ampC_0094 | blaCMY-2 | 32 | 38 | D | | | | | | | | | | | | | | | | | | | | | | | | | | |
| ampC_0095 | blaCMY-2 | 12 | 429 | B2 | | | | A | | | | | | | | | | | | | | | | | | | | | | |
| ampC_0096 | blaCMY-2 | 8 | 117 | F | | | | | | | | | | | | | | | | | | | | | | | | | | |
| ampC_0097 | blaCMY-2 | 12 | 429 | B2 | | | | A | | | | | | | | | | | | | | | | | | | | | | |
| ampC_0098 | blaCMY-2 | 3 | 1196 | B1 | | | | | | | | A | | | | | | | | | | | | | | | | | | |
| ampC_0099 | blaCMY-2 | 8 | 38 | D | | | | | | | | | | | | | | | | | | | | | | | | | | |
| ampC_0100 | blaCMY-2 | 6 | 88 | C | | | | | | | | A | | | | | | | | | | | | | | | | | | |
| ampC_0101 | blaCMY-2 | 8 | 38 | D | | | | | | | | | | | | | | | | | | | | | | | | | | |
| ampC_0102 | blaCMY-2 | 3 | 1196 | B1 | | | | | | | | A | | | | | | | | | | | | | | | | | | |
| ampC_0103 | blaCMY-2 | 8 | 38 | D | | | | | | | | | | | | | | | | | | | | | | | | | | |
| ampC_0104 | blaCMY-2 | 32 | 1485 | F | | | | A | | | | | | | | | | | | | | | | | | | | | | |
| ampC_0105 | blaCMY-2 | 6 | 429 | B2 | | | | A | | | | | | | | | | | | | | | | | | | | | | |
| ampC_0106 | blaCMY-2 | 8 | 101 | B1 | | | | | | | | A | | | | | | | | | | | | | | | | | | |
| ampC_0107 | blaCMY-2 | 8 | 38 | D | | | | | | | | | | | | | | | | | | | | | | | | | | |
| ampC_0108 | blaCMY-2 | 12 | 711 | B1 | | | | | | | | A | | | | | | | | | | | | | | | | | | |
| ampC_0109 | blaCMY-2 | 6 | unknown ST | A | | | | | | | | | | | | | | | | | | | | | | | | | | |
| ampC_0110 | blaCMY-2 | 8 | 429 | B2 | | | | A | | | | | | | | | | | | | | | | | | | | | | |
| ampC_0111 | blaCMY-2 | 12 | 118 | E | | | | | | | | | | | | | | | | | | | | | | | | | | |
| **ampC_0112** | **low-level** | **3** | **101** | **B1** | | | | | | G | | A | T | | | | | | | | | | | | | | | | | |
| ampC_0113 | blaDHA-1 | 1.5 | 394 | D | | | | | | | | | | | | | | | | | | | | | A | | | | | |
| ampC_0114 | blaCMY-2 | 32 | 5150 | D | | | | | | | | | | | | | | | | | | | | | | | | | | |
| ampC_0115 | hyperproducer | 0.75 | 131 | B2 | | | A | A | | | | | | | | | | | | | | | | | | | | | | |
| ampC_0116 | blaCMY-2 | 6 | 69 | D | | | | | | | | | | | | | | | | | | | | | | | | | | |
| ampC_0117 | blaCMY-2 | 8 | 38 | D | | | | | | | | | | | | | | | | | | | | | | | | | | |
| **ampC_0118** | **hyperproducer** | **4** | **95** | **B2** | | | | A | | | | | GT | | | | T | | | | | | | | | | | | | |
| ampC_0119 | blaCMY-4 & blaDHA-1 | 24 | 648 | F | | | | A | | | | | | | | | | | | | | | | | | | | A | | |
| ampC_0120 | blaCMY-2 | 32 | 973 | D | | | | | | | | | | | | | | | T | | | G | T | | | A | | | | |
| ampC_0121 | hyperproducer | 1.5 | 38 | D | | | A | | | | | | | | | | | | | | | | | | | | | | | |
| ampC_0122 | blaDHA-1 | 1 | 34 | A | | | | | | | | | | | | | | | | | | | | | | | | | | |
| ampC_0123 | blaCMY-2 | 12 | 69 | D | | | | | | | | | | | | | | | | | | | | | | | | | | |
| ampC_0124 | hyperproducer | 1.5 | 200 | B1 | | T | | | | | | A | | | | | | | | | | | | | | | | | | |
| ampC_0125 | hyperproducer | 1 | unknown ST | B2 | | | A | | | | | | | | | | | | | | | | | | | | | | | |
| ampC_0126 | low-level | 0.75 | 135 | B2 | | | | A | | | | | | | | | T | | | | | | | | | | | | | |
| ampC_0127 | low-level | 0.75 | unknown ST | D | | | | | | | C | | | | | T | | | | | | | | | | | | | | |
| **ampC_0128** | **hyperproducer** | **24** | **5358** | **cladeIV** | del ATCC | | | | | | | A | | | | | | G | | | | A | T | A | | | | | | |
| **ampC_0129** | **hyperproducer** | **3** | **73** | **B2** | | | | | | | | | | T | | | | | | | | | | | | | | | | |
| ampC_0130 | low-level | 0.38 | unknown ST | B2 | | | | A | | | T | | | | | T | T | | | | | | | | | | | | | |
| ampC_0131 | blaCMY-2 | 32 | 131 | B2 | | | | A | | | | | | | | | | | | | | | | | | | | | | |
| ampC_0132 | hyperproducer | 1 | 707 | B1 | | T | | | | | | A | | | | | | | | | | | | | | | | | | |
| **ampC_0133** | **hyperproducer** | **3** | **1423** | **B1** | | T | | | | | | A | | | | | | | | | | | | | | | | | | |
| **ampC_0134** | **hyperproducer** | **3** | **2015** | **B2** | | | A | | | | | | | | | | | | | A | | | | | | | | | | |
| **ampC_0135** | **hyperproducer** | **4** | **unknown ST** | **B1** | | T | | | | | | A | | | | | | | | | | | | | | | | | | |
| **ampC_0136** | **hyperproducer** | **8** | **167** | **A** | | T | | | | | | A | | | | | | | | | | | | | | | | | | |
| **ampC_0137** | **hyperproducer** | **3** | **345** | **B1** | | T | | | | | | A | | | | | | | del | | | | | | | | | | | |
| **ampC_0138** | **hyperproducer** | **3** | **88** | **C** | | T | | | | | | A | | | | | | | | | A | | | | | | | | | |
| ampC_0139 | hyperproducer | 2 | unknown ST | B1 | | T | | | | | | A | | | | | | | | | | | | | | | | | | |
| ampC_0140 | hyperproducer | 1 | 95 | B2 | | | | A | | | | | | T | | | T | | | | | | | | | | | | | |
| ampC_0141 | hyperproducer | 0.38 | 141 | B2 | | | | | | | | | | G | | | | | | | | | | | | | | | | |
| ampC_0142 | hyperproducer | 0.75 | 141 | B2 | | | A | | | | | | | | | | | | | | | | | | | A | | | | |
| ampC_0143 | hyperproducer | 1.5 | 88 | C | | | A | | | | | A | | | | | | | | A | | | | | | | | | | |
| ampC_0144 | hyperproducer | 0.5 | 73 | B2 | | | A | | | | | | | | | | | | | | | | | | | | | | | |
| **ampC_0145** | **hyperproducer** | **4** | **88** | **C** | | T | | | | | | A | | | | | | | | | | | | | | | | | | |
| **ampC_0146** | **hyperproducer** | **3** | **12** | **B2** | | | A | | | | | | | | | | | | | | | | | | | | | | | |
| ampC_0147 | blaCMY-2 | 24 | 131 | B2 | | | | A | | | | | | | | | | | | | | | | | | | | | | |
| ampC_0148 | low-level | 0.5 | 95 | B2 | | | | A | | | | | C | | | | T | | | | | | | | | | | | | |
| ampC_0149 | hyperproducer | 0.5 | 131 | B2 | | | A | A | | | | | | | | | | | | | | | | | | | | | | |
| ampC_0150 | hyperproducer | 0.5 | 73 | B2 | | | | | | | | | | G | | | | | | | | | | | | | | | | |
| ampC_0151 | hyperproducer | 1.5 | 1057 | B2 | | | A | A | | | | | | | | | T | | | | | | | | | | | | | |
| ampC_0152 | hyperproducer | 1 | 2328 | B1 | | T | | | | | | A | | | | | | | | | | | | | | | | | | |
| ampC_0153 | low-level | 0.19 | 73 | B2 | | | | | | | T | | | | | | | | | | | | | | | | | | G | |
| ampC_0154 | hyperproducer | 1 | 394 | D | | | A | | | | | | | | | | | | | | | | | | A | | | | | |
| ampC_0155 | hyperproducer | 0.75 | 95 | B2 | | | A | A | | | | | | | | | T | | | | | | | | | | | | | |
| ampC_0156 | hyperproducer | 1.5 | 973 | D | | | | | | | | | | | T | | | | T | | | G | T | | | A | | | | |
| ampC_0157 | hyperproducer | 0.75 | 968 | B2 | | | A | | | | | | | | | | | | | | | | | | | | | | | |
| ampC_0158 | hyperproducer | 0.38 | 73 | B2 | | | A | | | | | | | | | | | | | | | | | | | | | | | |
| ampC_0159 | blaDHA-1 | 4 | 349 | D | | | | | | | | | | | | | | | | | | | | | | | | | | T |
| ampC_0160 | low-level | 0.75 | 390 | B2 | | | | A | T | | | T | | | | | T | | A | | | | | | | | | | | |
| **ampC_0161** | **hyperproducer** | **3** | **73** | **B2** | | | | | | | | | | | T | | | | | | | | | | | | | | | |
| ampC_0162 | hyperproducer | 1 | 12 | B2 | | | | | | | | | GT | | | | | | | | | | | | | | | | | |
| ampC_0163 | blaCMY-2 | 32 | 429 | B2 | | | | A | | | | | | | | | | | | | | | | | | | | | | |
| **ampC_0164** | **hyperproducer** | **4** | **8190** | **B1** | | T | | | | | | A | | | | | | | | | | | | | | | | | | |
| ampC_0165 | blaCMY-2 | 12 | 12 | B2 | | | | | | | | | | | | | | | | | | | | | | | | | | |
| **ampC_0166** | **hyperproducer** | **4** | **641** | **B1** | | T | | | | | | A | | | | | | | | | | | | | | | | | | |
| ampC_0167 | hyperproducer | 2 | 357 | B2 | | | | A | | | | | | | T | | | | A | | | | | | | | | | | |
| ampC_0168 | blaCMY-2 | 8 | 355 | B2 | | | | A | | | | | | | | | T | | | | | | | | | | | | | |
| ampC_0169 | blaCMY-2 | 32 | 453 | B1 | | | | | | | | | | | | | | | | | | | | | | | | | | |
| **ampC_0170** | **hyperproducer** | **32** | **99** | **B1** | | T | | | | | | A | | | | | | | | | | | | | | | A | | | |
| ampC_0171 | hyperproducer | 2 | 1674 | F | | T | | | | | | A | | | | | | | | | | | | | | | | | | |
| ampC_0172 | hyperproducer | 0.75 | 73 | B2 | | | | | | | | | | G | | | | | | | | | | | | | | | | |
