## Supplemental Table 2 for "Genome-wide analysis in *Escherichia coli* unravels an unprecedented level of genetic homoplasy associated with cefotaxime resistance"

**Table S2**

SNP analysis for *n*=172 *E. coli* isolates according to snippy statistics.

| | LENGTH | ALIGNED | UNALIGNED | VARIANT | HET[a] | MASKED | LOWCOV[b] |
|---|---|---|---|---|---|---|---|
| ampC_0001 | 5056572 | 4002058 | 1020001 | 91335 | 207 | 0 | 34306 |
| ampC_0002 | 5056572 | 4319724 | 714013 | 74758 | 3653 | 0 | 19182 |
| ampC_0003 | 5056572 | 4329143 | 711707 | 72187 | 2225 | 0 | 13497 |
| ampC_0004 | 5056572 | 4086651 | 946901 | 79975 | 3771 | 0 | 19249 |
| ampC_0005 | 5056572 | 4118550 | 908345 | 79937 | 1750 | 0 | 27927 |
| ampC_0006 | 5056572 | 4412142 | 620848 | 69875 | 2349 | 0 | 21233 |
| ampC_0007 | 5056572 | 4130043 | 899616 | 79049 | 3795 | 0 | 23118 |
| ampC_0008 | 5056572 | 4183807 | 851236 | 77310 | 2729 | 0 | 18800 |
| ampC_0009 | 5056572 | 4200448 | 829949 | 65966 | 1995 | 0 | 24180 |
| ampC_0010 | 5056572 | 4165334 | 863696 | 75495 | 2561 | 0 | 24981 |
| ampC_0011 | 5056572 | 4059076 | 981897 | 75567 | 2136 | 0 | 13463 |
| ampC_0012 | 5056572 | 4035669 | 964704 | 73761 | 1380 | 0 | 54819 |
| ampC_0013 | 5056572 | 4256665 | 775179 | 64897 | 3719 | 0 | 21009 |
| ampC_0014 | 5056572 | 4433064 | 602743 | 63312 | 4503 | 0 | 16262 |
| ampC_0015 | 5056572 | 3958952 | 1079669 | 59707 | 1017 | 0 | 16934 |
| ampC_0016 | 5056572 | 4097364 | 934798 | 59872 | 891 | 0 | 23519 |
| ampC_0017 | 5056572 | 4075287 | 963082 | 58030 | 1860 | 0 | 16343 |
| ampC_0018 | 5056572 | 4094996 | 944951 | 58091 | 1347 | 0 | 15278 |
| ampC_0019 | 5056572 | 3947082 | 873023 | 46255 | 153178 | 0 | 83289 |
| ampC_0020 | 5056572 | 4086689 | 941661 | 50545 | 839 | 0 | 27383 |
| ampC_0021 | 5056572 | 4139227 | 895700 | 48942 | 3323 | 0 | 18322 |
| ampC_0022 | 5056572 | 4574566 | 452688 | 13190 | 1247 | 0 | 28071 |
| ampC_0023 | 5056572 | 4082302 | 948985 | 47881 | 2458 | 0 | 22827 |
| ampC_0024 | 5056572 | 4188724 | 844580 | 48264 | 1491 | 0 | 21777 |
| ampC_0025 | 5056572 | 4117554 | 922183 | 49263 | 1624 | 0 | 15211 |
| ampC_0026 | 5056572 | 4352434 | 689821 | 30020 | 3894 | 0 | 10423 |
| ampC_0027 | 5056572 | 4421498 | 595332 | 42789 | 3204 | 0 | 36538 |
| ampC_0028 | 5056572 | 4075024 | 953944 | 49287 | 2526 | 0 | 25078 |
| ampC_0029 | 5056572 | 4269805 | 769936 | 44014 | 2395 | 0 | 14436 |
| ampC_0030 | 5056572 | 4566336 | 448506 | 13088 | 1005 | 0 | 40725 |
| ampC_0031 | 5056572 | 4120182 | 923997 | 49065 | 1475 | 0 | 10918 |
| ampC_0032 | 5056572 | 4134701 | 897045 | 49046 | 1696 | 0 | 23130 |
| ampC_0033 | 5056572 | 4322514 | 711940 | 43498 | 2754 | 0 | 19364 |
| ampC_0034 | 5056572 | 4174080 | 860015 | 43604 | 1241 | 0 | 21236 |
| ampC_0035 | 5056572 | 4224196 | 798976 | 47616 | 1862 | 0 | 31538 |

| | | | | | | | |
|---|---|---|---|---|---|---|---|
| ampC_0036 | 5056572 | 4090346 | 952537 | 47418 | 977 | 0 | 12712 |
| ampC_0037 | 5056572 | 4122180 | 897623 | 46351 | 1069 | 0 | 35700 |
| ampC_0038 | 5056572 | 4030895 | 1007575 | 48994 | 926 | 0 | 17176 |
| ampC_0039 | 5056572 | 4062961 | 976111 | 48679 | 4308 | 0 | 13192 |
| ampC_0040 | 5056572 | 4043013 | 998179 | 49855 | 1057 | 0 | 14323 |
| ampC_0041 | 5056572 | 4185762 | 838606 | 47709 | 2353 | 0 | 29851 |
| ampC_0042 | 5056572 | 4103666 | 934960 | 48932 | 3264 | 0 | 14682 |
| ampC_0043 | 5056572 | 4015738 | 1020522 | 49034 | 1587 | 0 | 18725 |
| ampC_0044 | 5056572 | 4739734 | 304443 | 3813 | 2772 | 0 | 9623 |
| ampC_0045 | 5056572 | 4112373 | 923560 | 49142 | 1125 | 0 | 19514 |
| ampC_0046 | 5056572 | 4181283 | 858993 | 46814 | 1130 | 0 | 15166 |
| ampC_0047 | 5056572 | 4099314 | 942142 | 47041 | 353 | 0 | 14763 |
| ampC_0048 | 5056572 | 4148746 | 888010 | 46907 | 2702 | 0 | 17114 |
| ampC_0049 | 5056572 | 4262477 | 776751 | 49015 | 1870 | 0 | 15474 |
| ampC_0050 | 5056572 | 4239904 | 695026 | 41758 | 1797 | 0 | 119845 |
| ampC_0051 | 5056572 | 4237499 | 798907 | 46047 | 2417 | 0 | 17749 |
| ampC_0052 | 5056572 | 4213115 | 827100 | 46186 | 2238 | 0 | 14119 |
| ampC_0053 | 5056572 | 4222395 | 815897 | 42407 | 1832 | 0 | 16448 |
| ampC_0054 | 5056572 | 4377423 | 657887 | 28917 | 4262 | 0 | 17000 |
| ampC_0055 | 5056572 | 4099407 | 939857 | 45724 | 597 | 0 | 16711 |
| ampC_0056 | 5056572 | 4204631 | 833416 | 43095 | 1140 | 0 | 17385 |
| ampC_0057 | 5056572 | 4154961 | 880433 | 47379 | 3120 | 0 | 18058 |
| ampC_0058 | 5056572 | 4186917 | 788884 | 41961 | 802 | 0 | 79969 |
| ampC_0059 | 5056572 | 4070307 | 963077 | 46192 | 1370 | 0 | 21818 |
| ampC_0060 | 5056572 | 4075004 | 960143 | 47202 | 2713 | 0 | 18712 |
| ampC_0061 | 5056572 | 4131932 | 898004 | 45626 | 1687 | 0 | 24949 |
| ampC_0062 | 5056572 | 4049499 | 985262 | 48089 | 2511 | 0 | 19300 |
| ampC_0063 | 5056572 | 4142668 | 895138 | 44464 | 2335 | 0 | 16431 |
| ampC_0064 | 5056572 | 4093514 | 945558 | 45406 | 1576 | 0 | 15924 |
| ampC_0065 | 5056572 | 4052337 | 946163 | 46789 | 2526 | 0 | 55546 |
| ampC_0066 | 5056572 | 4309198 | 698918 | 41471 | 2051 | 0 | 46405 |
| ampC_0067 | 5056572 | 4098125 | 930908 | 47362 | 2095 | 0 | 25444 |
| ampC_0068 | 5056572 | 4034970 | 998701 | 47682 | 974 | 0 | 21927 |
| ampC_0069 | 5056572 | 4963049 | 88082 | 0 | 2126 | 0 | 3315 |
| ampC_0070 | 5056572 | 4261402 | 758981 | 41647 | 2298 | 0 | 33891 |
| ampC_0071 | 5056572 | 4062083 | 979818 | 47899 | 1675 | 0 | 12996 |
| ampC_0072 | 5056572 | 4122405 | 919634 | 46230 | 1606 | 0 | 12927 |
| ampC_0073 | 5056572 | 4083201 | 960580 | 47439 | 2455 | 0 | 10336 |
| ampC_0074 | 5056572 | 4261912 | 776279 | 42020 | 4323 | 0 | 14058 |
| ampC_0075 | 5056572 | 4141662 | 898054 | 45227 | 3219 | 0 | 13637 |

| | | | | | | | |
|---|---|---|---|---|---|---|---|
| ampC_0076 | 5056572 | 4406231 | 637952 | 40756 | 2070 | 0 | 10319 |
| ampC_0077 | 5056572 | 4074836 | 966420 | 47889 | 2057 | 0 | 13259 |
| ampC_0078 | 5056572 | 4136130 | 899450 | 45683 | 3415 | 0 | 17577 |
| ampC_0079 | 5056572 | 4075563 | 960304 | 46693 | 2343 | 0 | 18362 |
| ampC_0080 | 5056572 | 4145656 | 887683 | 42153 | 3022 | 0 | 20211 |
| ampC_0081 | 5056572 | 4401616 | 639375 | 40756 | 2160 | 0 | 13421 |
| ampC_0082 | 5056572 | 4171044 | 866701 | 45273 | 2024 | 0 | 16803 |
| ampC_0083 | 5056572 | 4068141 | 967971 | 47390 | 1834 | 0 | 18626 |
| ampC_0084 | 5056572 | 4061000 | 967098 | 47391 | 1649 | 0 | 26825 |
| ampC_0085 | 5056572 | 4088462 | 949229 | 47379 | 1884 | 0 | 16997 |
| ampC_0086 | 5056572 | 4275337 | 756692 | 41604 | 3109 | 0 | 21434 |
| ampC_0087 | 5056572 | 4386513 | 652348 | 41247 | 3261 | 0 | 14450 |
| ampC_0088 | 5056572 | 4318192 | 701346 | 41226 | 4400 | 0 | 32634 |
| ampC_0089 | 5056572 | 4023554 | 1017880 | 46064 | 1504 | 0 | 13634 |
| ampC_0090 | 5056572 | 4128921 | 903372 | 43885 | 1087 | 0 | 23192 |
| ampC_0091 | 5056572 | 4157584 | 876159 | 43838 | 1302 | 0 | 21527 |
| ampC_0092 | 5056572 | 4259206 | 778237 | 47245 | 3166 | 0 | 15963 |
| ampC_0093 | 5056572 | 4052051 | 983719 | 47518 | 1770 | 0 | 19032 |
| ampC_0094 | 5056572 | 4222190 | 815495 | 40990 | 1463 | 0 | 17424 |
| ampC_0095 | 5056572 | 4046770 | 985438 | 47427 | 1735 | 0 | 22629 |
| ampC_0096 | 5056572 | 4149354 | 867928 | 41561 | 4625 | 0 | 34665 |
| ampC_0097 | 5056572 | 4046706 | 984650 | 47430 | 1697 | 0 | 23519 |
| ampC_0098 | 5056572 | 4128686 | 901637 | 44668 | 1477 | 0 | 24772 |
| ampC_0099 | 5056572 | 4220835 | 810119 | 40979 | 1539 | 0 | 24079 |
| ampC_0100 | 5056572 | 4239640 | 782839 | 44934 | 1806 | 0 | 32287 |
| ampC_0101 | 5056572 | 4222079 | 813239 | 40990 | 1515 | 0 | 19739 |
| ampC_0102 | 5056572 | 4110515 | 922832 | 44646 | 867 | 0 | 22358 |
| ampC_0103 | 5056572 | 4224370 | 811359 | 40935 | 1931 | 0 | 18912 |
| ampC_0104 | 5056572 | 4497404 | 545801 | 12830 | 2198 | 0 | 11169 |
| ampC_0105 | 5056572 | 4053473 | 981533 | 47259 | 1937 | 0 | 19629 |
| ampC_0106 | 5056572 | 4106292 | 931548 | 44164 | 1320 | 0 | 17412 |
| ampC_0107 | 5056572 | 4226657 | 810011 | 40862 | 1873 | 0 | 18031 |
| ampC_0108 | 5056572 | 4117735 | 921267 | 44203 | 1431 | 0 | 16139 |
| ampC_0109 | 5056572 | 4057561 | 982568 | 45389 | 1350 | 0 | 15093 |
| ampC_0110 | 5056572 | 4048316 | 987588 | 46969 | 2042 | 0 | 18626 |
| ampC_0111 | 5056572 | 4226287 | 804856 | 46319 | 2887 | 0 | 22542 |
| ampC_0112 | 5056572 | 4091890 | 944221 | 74707 | 1289 | 0 | 19172 |
| ampC_0113 | 5056572 | 4327109 | 710270 | 65760 | 3513 | 0 | 15680 |
| ampC_0114 | 5056572 | 4369682 | 670026 | 64621 | 3765 | 0 | 13099 |
| ampC_0115 | 5056572 | 4099316 | 940214 | 73818 | 1516 | 0 | 15526 |

| | | | | | | | |
|---|---|---|---|---|---|---|---|
| ampC_0116 | 5056572 | 4250258 | 785532 | 64763 | 1541 | 0 | 19241 |
| ampC_0117 | 5056572 | 4419273 | 617244 | 40771 | 3377 | 0 | 16678 |
| ampC_0118 | 5056572 | 4073836 | 944236 | 46719 | 1746 | 0 | 36754 |
| ampC_0119 | 5056572 | 4767553 | 281363 | 868 | 1448 | 0 | 6208 |
| ampC_0120 | 5056572 | 4178761 | 857249 | 41249 | 714 | 0 | 19848 |
| ampC_0121 | 5056572 | 4417409 | 617358 | 40872 | 3643 | 0 | 18162 |
| ampC_0122 | 5056572 | 4154418 | 882714 | 45535 | 2680 | 0 | 16760 |
| ampC_0123 | 5056572 | 4282639 | 748368 | 40715 | 3251 | 0 | 22314 |
| ampC_0124 | 5056572 | 4199402 | 833286 | 44226 | 3584 | 0 | 20300 |
| ampC_0125 | 5056572 | 4071153 | 939352 | 45824 | 1277 | 0 | 44790 |
| ampC_0126 | 5056572 | 4057481 | 967449 | 72037 | 535 | 0 | 31107 |
| ampC_0127 | 5056572 | 4207961 | 792907 | 65427 | 1733 | 0 | 53971 |
| ampC_0128 | 5056572 | 3591028 | 1361797 | 208032 | 502 | 0 | 103245 |
| ampC_0129 | 5056572 | 4075848 | 935972 | 65680 | 2703 | 0 | 42049 |
| ampC_0130 | 5056572 | 4078609 | 901698 | 65163 | 1614 | 0 | 74651 |
| ampC_0131 | 5056572 | 4015457 | 987193 | 65686 | 1309 | 0 | 52613 |
| ampC_0132 | 5056572 | 4075070 | 932609 | 65329 | 606 | 0 | 48287 |
| ampC_0133 | 5056572 | 4086975 | 913964 | 65140 | 498 | 0 | 55135 |
| ampC_0134 | 5056572 | 4074172 | 926688 | 63703 | 1428 | 0 | 54284 |
| ampC_0135 | 5056572 | 4128412 | 879533 | 65081 | 1125 | 0 | 47502 |
| ampC_0136 | 5056572 | 4133358 | 863522 | 65141 | 2064 | 0 | 57628 |
| ampC_0137 | 5056572 | 4112205 | 888398 | 64554 | 843 | 0 | 55126 |
| ampC_0138 | 5056572 | 4192841 | 796407 | 64562 | 1743 | 0 | 65581 |
| ampC_0139 | 5056572 | 4123583 | 893959 | 64542 | 639 | 0 | 38391 |
| ampC_0140 | 5056572 | 4175232 | 810119 | 62832 | 2496 | 0 | 68725 |
| ampC_0141 | 5056572 | 4063097 | 939756 | 63842 | 1609 | 0 | 52110 |
| ampC_0142 | 5056572 | 4043055 | 961717 | 63808 | 1908 | 0 | 49892 |
| ampC_0143 | 5056572 | 4252968 | 757696 | 64399 | 1304 | 0 | 44604 |
| ampC_0144 | 5056572 | 4050534 | 941204 | 63302 | 2995 | 0 | 61839 |
| ampC_0145 | 5056572 | 4189356 | 797298 | 44281 | 819 | 0 | 69099 |
| ampC_0146 | 5056572 | 4042816 | 955684 | 63450 | 1298 | 0 | 56774 |
| ampC_0147 | 5056572 | 4048924 | 965151 | 64008 | 1492 | 0 | 41005 |
| ampC_0148 | 5056572 | 4030869 | 924256 | 62459 | 2116 | 0 | 99331 |
| ampC_0149 | 5056572 | 4102015 | 908081 | 62696 | 1112 | 0 | 45364 |
| ampC_0150 | 5056572 | 4061005 | 939691 | 62730 | 2972 | 0 | 52904 |
| ampC_0151 | 5056572 | 4072426 | 928748 | 62427 | 2780 | 0 | 52618 |
| ampC_0152 | 5056572 | 4120065 | 870133 | 64191 | 1144 | 0 | 65230 |
| ampC_0153 | 5056572 | 4073460 | 927811 | 62762 | 3559 | 0 | 51742 |
| ampC_0154 | 5056572 | 4276093 | 716808 | 56622 | 2590 | 0 | 61081 |
| ampC_0155 | 5056572 | 4201782 | 796742 | 61827 | 2586 | 0 | 55462 |

| | | | | | | | |
|---|---|---|---|---|---|---|---|
| ampC_0156 | 5056572 | 4190874 | 819893 | 56296 | 1403 | 0 | 44402 |
| ampC_0157 | 5056572 | 4031908 | 966021 | 45951 | 2745 | 0 | 55898 |
| ampC_0158 | 5056572 | 4019423 | 982398 | 62085 | 2279 | 0 | 52472 |
| ampC_0159 | 5056572 | 4226429 | 763993 | 57038 | 1274 | 0 | 64876 |
| ampC_0160 | 5056572 | 4081580 | 910039 | 61699 | 1862 | 0 | 63091 |
| ampC_0161 | 5056572 | 4060774 | 951880 | 61950 | 1797 | 0 | 42121 |
| ampC_0162 | 5056572 | 3901480 | 953803 | 58461 | 1335 | 0 | 199954 |
| ampC_0163 | 5056572 | 4057724 | 945684 | 60127 | 2385 | 0 | 50779 |
| ampC_0164 | 5056572 | 4157175 | 841316 | 60662 | 988 | 0 | 57093 |
| ampC_0165 | 5056572 | 4030168 | 971013 | 61326 | 2923 | 0 | 52468 |
| ampC_0166 | 5056572 | 4068330 | 935049 | 60310 | 808 | 0 | 52385 |
| ampC_0167 | 5056572 | 4061853 | 946454 | 60045 | 680 | 0 | 47585 |
| ampC_0168 | 5056572 | 4004238 | 981827 | 58506 | 1569 | 0 | 68938 |
| ampC_0169 | 5056572 | 4144719 | 858069 | 60377 | 982 | 0 | 52802 |
| ampC_0170 | 5056572 | 4137458 | 861197 | 60084 | 773 | 0 | 57144 |
| ampC_0171 | 5056572 | 4342878 | 684949 | 36317 | 423 | 0 | 28322 |
| ampC_0172 | 5056572 | 4065467 | 932423 | 58567 | 3198 | 0 | 55484 |

[a]Heterozygous variants, [b]SNPs with coverage <20-fold
